# Corticothalamic feedback sculpts visual spatial integration in mouse thalamus

**DOI:** 10.1101/2020.05.19.104000

**Authors:** Gregory Born, Felix A. Schneider, Sinem Erisken, Agne Klein, Chu Lan Lao, Milad H. Mobarhan, Martin A. Spacek, Gaute T. Einevoll, Laura Busse

## Abstract

En route from retina to cortex, visual information passes through the dorsolateral geniculate nucleus of the thalamus (dLGN), where extensive corticothalamic (CT) feedback has been suggested to modulate spatial processing. How this modulation arises from direct excitatory and indirect inhibitory CT feedback pathways remains enigmatic. Here we show that in awake mice, retinotopically organized cortical feedback sharpens receptive fields (RFs) and increases surround suppression in the dLGN. Guided by a network model indicating that widespread inhibitory CT feedback is necessary to reproduce these effects, we targeted the visual sector of the thalamic reticular nucleus (visTRN) for recordings. We found that visTRN neurons have large receptive fields, show little surround suppression, and exhibit strong feedback-dependent responses to large stimuli. These features make them an ideal candidate for mediating feedback-enhanced surround suppression in the dLGN. We conclude that cortical feedback sculpts spatial integration in dLGN, likely via recruitment of neurons in visTRN.

## Introduction

According to the classical view of the visual system, information is propagated through a hierarchy of processing stages, interconnected via feedforward projections. The initial feedforward sweep can contain a significant amount of information, sometimes sufficient to drive perception^1–3^. Furthermore, feedforward architectures implemented in artificial neural networks can give rise to representations that resemble those in the primate visual processing hierarchy, enabling human-like performance in complex visual tasks, such as object recognition^4, 5^.

While feedforward architectures can be powerful, anatomical connectivity in the brain is dominated by feedback. Feedback is a prominent and ubiquitous motif in the brain, observed across areas and species, with descending feedback projections generally outnumbering ascending feedforward afferents. For instance, most inputs to primary sensory cortical areas do not come from the primary thalamus, but from higher-order regions^6^. The same principle applies to corticothalamic (CT) communication: relay cells in the dorsolateral geniculate nucleus (dLGN) of the thalamus receive only 5-10% of their synaptic inputs from retinal afferents, whereas 30% originate from L6 corticothalamic (L6CT) pyramidal cells in the primary visual cortex (V1)^7^. While this pronounced prevalence implies that feedback projections provide core aspects of neural computation, a consensus regarding the function of feedback to lower-level sensory areas is lacking^8, 9^.

Largely conserved across mammalian species and similarly organized across the visual, auditory, and somatosensory systems^10–12^, the CT feedback circuit is ideal for investigating the role of feedback; yet how CT feedback influences the representation of information in thalamus remains poorly understood. In the visual system, for instance, despite massive cortical input, dLGN RFs more closely resemble retinal than cortical RFs^13–16^. Indeed, instead of providing driving input, CT feedback is considered to be a modulator of thalamic responses, and is implicated in more subtle changes in thalamic RFs^17^. These alterations will be complex, because L6CT pyramidal cells not only provide direct excitation to thalamic relay cells, but also indirect inhibition, by exciting inhibitory neurons in the thalamic reticular nucleus (TRN) and local dLGN interneurons. The specific balance of feedback-mediated excitation and indirect inhibition is likely dynamic and flexible, given the distinct stimulus selectivity and synaptic properties in the CT circuit^18, 19^.

Most notably, the retinotopic arrangement of CT projections in the visual system of primates and cats^20–24^ is highly suggestive of a role in modulating spatial processing, yet experiments on spatial integration have yielded conflicting results: while some argue that suppressive influences elicited by stimuli placed outside the RF of dLGN neurons rely entirely on intrathalamic or retinal mechanisms^25, 26^, other evidence suggests that CT feedback can enhance surround suppression^27–33^. Recent optogenetic investigations in mice have failed to clarify the matter. Indeed, while Olsen et al.^34^ report data consistent with a role of feedback in strengthening contextual influences of spatial processing, other studies have failed to observe any effect of CT feedback on spatial integration^35^, or indeed on any other aspect of mouse dLGN activity^36, 37^.

Here, we studied the role of cortical feedback in modulating thalamic spatial integration across the main processing stages of the thalamo-cortico-thalamic loop - V1, dLGN, and visual TRN (visTRN). Using viral labeling and channelrhodopsin (ChR2)-assisted functional mapping, we found that V1 corticogeniculate feedback projections in the mouse are retinotopically organized and have spatially specific function. Using optogenetic manipulations to compare the modulation by CT feedback of dLGN responses to stimuli of various sizes, we revealed that CT feedback sharpens RFs and increases surround suppression. Computational modelling and recordings from the TRN provided evidence that CT feedback can augment dLGN surround suppression via the visTRN. We conclude that CT feedback sculpts spatial integration in the dLGN, most probably via recruitment of inhibitory neurons in the TRN.

## Results

### Corticogeniculate feedback is retinotopic and exerts spatially specific effects

Modulations of spatial processing could be achieved efficiently if CT feedback in the mouse visual system was arranged in a retinotopic way. To test whether CT feedback terminal fields maintained the retinotopic organization from their V1 origin, we performed triple-color cre-dependent viral tracing, by injecting small volumes of AAV at different retinotopic positions into V1 of Ntsr1-Cre mice^38^, targeting with *>* 90% specificity L6CT pyramidal cells (**Figure 1a,b**). Visual inspection of thalamic slices revealed that the anterogradely labeled axonal terminal fields of different colors were clearly separated in dLGN, matched the pattern of expression in V1, and were consistent with the retinotopic organization of the geniculate target region^39^ (**Figure 1c**, **Figure S1**). Such spatially precise, retinotopic organization, as known from previous work in primates and cats^20–24^, make CT feedback projections well poised to modulate spatial representations.

**Figure 1.**
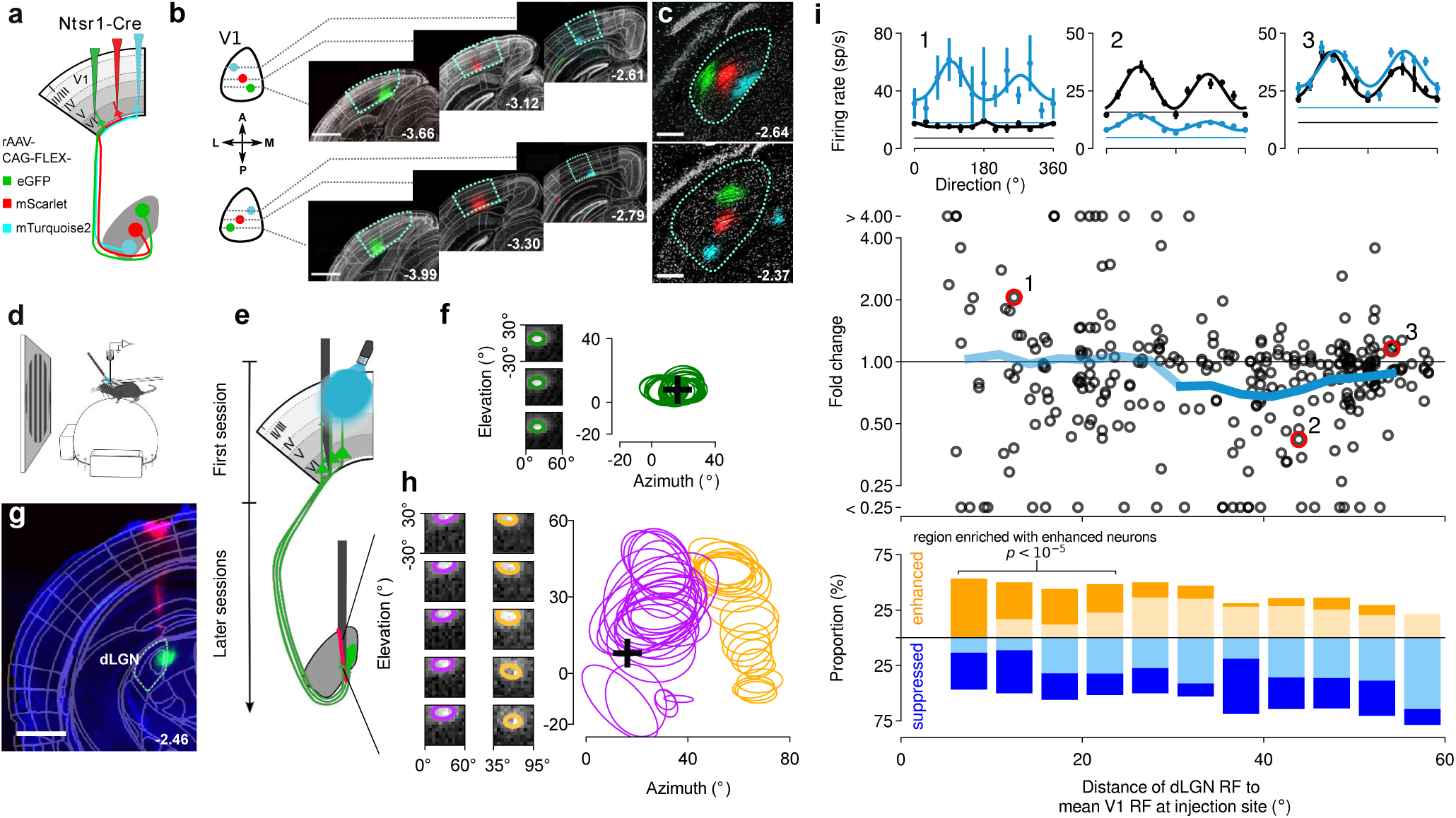
Anatomical and functional mapping of L6CT feedback. (**a**) Schematic of the triple-color viral tracing experiment, in which localized populations of V1 L6CT neurons were transduced with Cre-dependent AAV-FLEX-eGFP, AAV-FLEX-mScarlet, AAV-FLEX-mTurquoise2 in Ntsr1-Cre mice^38^. (**b**) Coronal sections of V1 showing the locations and extent of the injections along the V1 axis for azimuth (*top*) and elevation (*bottom*). *Green*: eGFP; *red*: mScarlet; *blue*: mTurquoise2. *Gray*: Nissl; scale bar 0.5 mm. Narrow-field images of *mTurquoise2* were acquired with a confocal microscope and manually aligned with the wide-field epifluorescence images of the corresponding brain slices. (**c**) Labeled V1 L6CT axonal terminal fields in dLGN, arranged in clear topographic order and consistent with the known retinotopy in dLGN^39^ (*top*: elevation axis; *bottom*: azimuth axis). (**d**) Schematic of setup for *in vivo* recordings in head-fixed mice. (**e**) Schematic of ChR2-assisted, functional connectivity mapping. (**f**) RF mapping in V1. *Left*: spatial RFs for three representative channels recorded at the V1 injection site. Colored circles represent 1 *σ* contours of the fitted 2D Gaussian. *Right*: all fitted V1 RF contours from the recording session shown. Black cross: mean RF center. (**g**) Coronal section showing local axonal termination of V1 L6CT neurons expressing ChR2-eYFP in dLGN. *Magenta*: DiI used to trace the recording track; Ileft: scale bar 1 mm. (**h**) RF mapping in dLGN. *Left*: spatial RFs and fitted 1 *σ* contours for representative channels located in the dLGN (two recording sessions, channel order top-to-bottom, as depicted on the probe in (e)). *Right*: fitted dLGN RF contours with mean RF position from the V1 recording indicated by the black cross. (**i**) Spatial profile of modulations induced by photostimulation of CT feedback. *Top*: direction-tuning curves for three neurons sampled during photostimulation of L6CT pyramidal cells (*blue*) and under control conditions (*black*); full-screen sine-wave gratings drifting in one of 12 directions, with temporal and spatial frequencies coarsely optimized for the recording, were presented for 0.75 s; in half of the trials, randomly interleaved, the stimulus was paired with continuous photostimulation, starting 0.085 s before stimulus onset and lasting for 0.85 s. *Middle*: CT feedback modulation strength (fold change) plotted as a function of retinotopic distance to the mean V1 RF position at the injection site. The *red circles* mark the neurons for which data are shown in the *upper panel*. The *blue line* indicates the mean of fold-change values in overlapping bins (bin size 15 deg, spacing 3.3 deg), and its *thick portion* delineates the region with significant mean fold change (n = 293 neurons, *p* = 6.8 x 10*^−^*^3^, cluster permutation test).*Bottom*: proportions of significantly enhanced (*dark orange*), suppressed (*dark blue*) and unmodulated neurons (*pale*).

To examine whether the retinotopically organized CT feedback can also generate spatially specific functional effects, we next probed the localized impact of CT feedback on dLGN responses using ChR2-assisted functional mapping (**Figure 1d,e**, **Figure S2**). We expressed ChR2-eYFP in a localized population of L6CT pyramidal cells by injecting a small volume of Cre-dependent AAV expressing ChR2-eYFP into V1 of Ntsr1-Cre mice^38^. Post-mortem, we confirmed the spatial specificity of transduction by registering the histological data from individual mice into a standardized 3D anatomical coordinate system (Allen Mouse Common Coordinate Framework [CCF]^40^) and extracting relative expression volumes in both V1 L6 and dLGN (**Figure S2**).

After sufficient time for expression, we performed silicon-probe recordings in head-fixed mice (**Figure 1d,e**). We first targeted the V1 injection site and used a sparse noise stimulus to estimate average RF location of the neurons in the V1 column of the transduced L6CT cells (**Figure 1f**). We then turned to the dLGN (**Figure 1g**), where RF mapping revealed a smooth retinotopic progression^39^, with RFs covering positions from upper to lower visual field for consecutive recording channels along the dorsoventral axis (**Figure 1h**). Multiple sessions with different insertions enabled us to measure dLGN RFs located at a wide range of distances from the averaged RF at the V1 injection site.

We then functionally mapped the spatial profile of CT feedback effects by photostimulating the local population of transduced L6CT pyramidal cells during the presentation of full-screen drifting gratings (**Figure 1i**). To avoid potentially confounding, state-dependent response-modulations^41, 42^, we only considered trials in which the animal was quiescent (speed *≤* 0.25 cm/s for *≥* 80% of the trial) for the computation of direction-tuning curves and thus for the evaluation of CT feedback effects. From these curves (**Figure 1i**, top), we determined, for each neuron, the relative CT feedback modulation strength as the ratio of responses under L6CT photostimulation to those seen under control conditions (fold change). Across the population of recorded dLGN neurons, activating CT feedback resulted in both enhancement (*n* = 112) and suppression of neuronal responses (*n* = 181), with diverse effect sizes. The diversity of effects was not obviously related to several aspects of tuning in the recorded dLGN neurons, such as recording depth, orientation selectivity or contrast sensitivity (**Figure S3**).

Plotting the responses of individual neurons to CT feedback modulation against their retinotopic distances from the activated L6CT pyramidal cell population, however, revealed a distinct spatial profile (**Figure 1i**, middle). While CT feedback, on average, had a small overall effect in retinotopically “nearby” regions (*<* 30 deg), average CT feedback modulation in “distant” regions was predominantly suppressive (30 *−* 53 deg, *p* = 6.8 *×* 10*^−^*^3^, cluster permutation test; thick blue line). The small average effect in “nearby” regions, rather than indicating the absence of modulation, reflected the diversity of modulatory effects (**Figure 1i**, bottom). Indeed, when we classified neurons into significantly enhanced, suppressed or not modulated by CT feedback, we observed that the prevalence of modulation types depended on retinotopic distance (*p* = 7.2 *×* 10*^−^*^3^, Chi-square test). Unlike suppressed neurons, which were uniformly distributed across retinotopic distances (*p* = 0.43, Chi-square test), numbers of enhanced neurons varied with distance (*p* = 4.7 *×* 10*^−^*^4^, Chi-square test) and were enriched in the sector from 0 *−* 25 deg (*p <* 10*^−^*^5^, cluster permutation test).

Together with our results of the triple-color viral tracing experiments, these findings demonstrate that L6CT pyramidal cell output impacts mouse dLGN in a spatially specific manner. The measured modulation profile is consistent with a circuit architecture in which enhancing influences of CT feedback are more tightly localized, while suppressive influences are dispersed over a wider spatial scale. This spatial profile, in particular the retinotopically distant suppressive region, is suggestive of L6CT feedback being involved in shaping dLGN spatial integration.

### CT feedback modulates dLGN spatial integration by sharpening RFs and increasing surround suppression

Having observed that photostimulation of CT feedback can, in principle, induce modulations of geniculate activity with a spatial profile suggestive of shaping spatial integration in dLGN, we set out to probe whether CT feedback is indeed involved in tuning for stimulus size and surround suppression (**Figure 2**). Surround suppression refers to the reduction of a neuron’s activity in response to a stimulus that exceeds the size of its classical RF (**Figure 2d**), and is thought to underlie fundamental aspects of visual processing, such as the integration and segregation of visual information^43, 44^ and the computation of perceptual salience^45, 46^. Furthermore, weakened center-surround interactions seem to be a low-level sensory facet of the compromised ability of schizophrenic patients to use context for the interpretation of stimuli^47, 48^.

**Figure 2.**
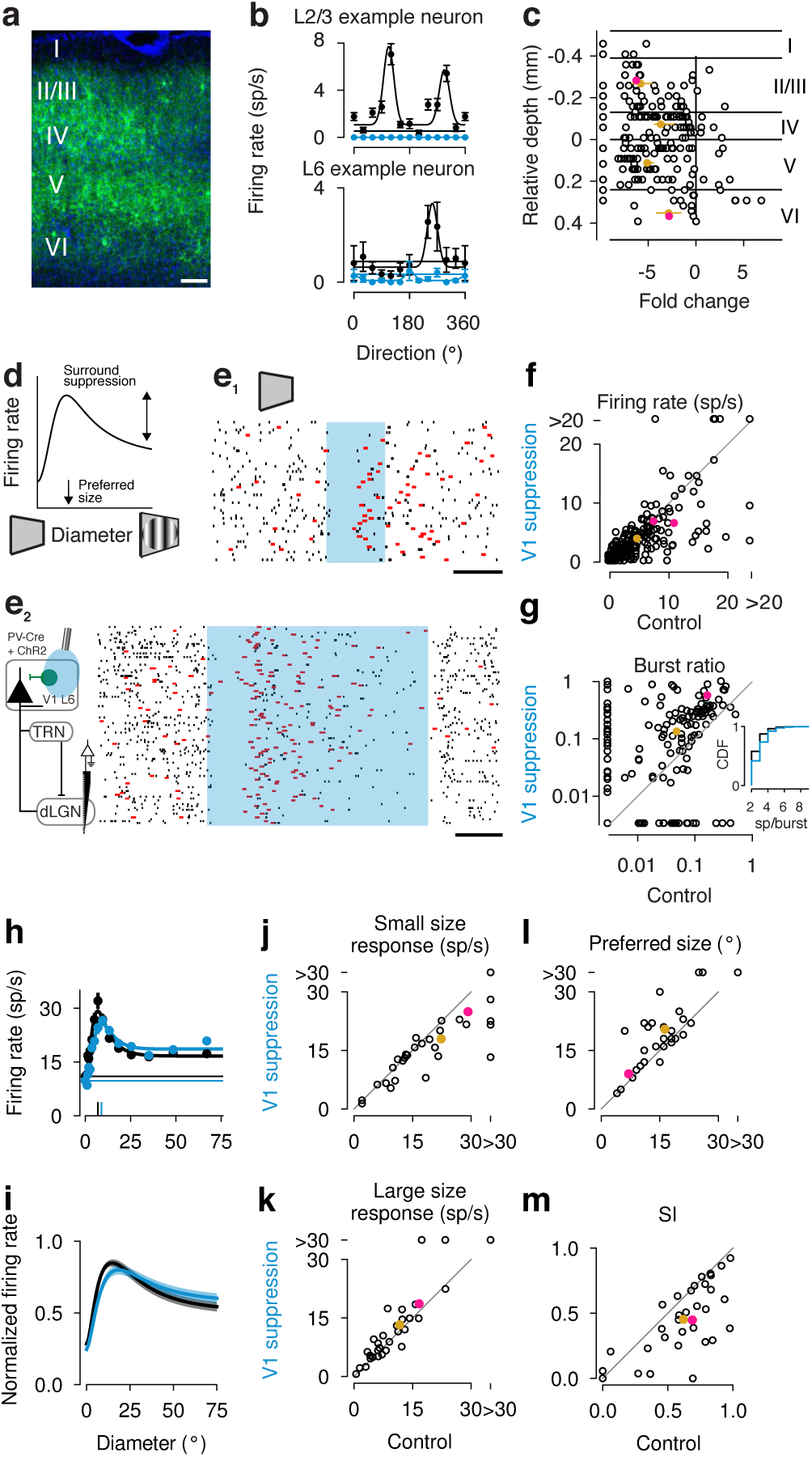
Suppression of cortical feedback modulates responses in dLGN in a stimulus-size-dependent manner. (**a**) Coronal section of V1 of a PV-Cre mouse injected with Cre-dependent AAV-ChR2. *Green*: ChR2-YFP, *blue*: DAPI. Scale bar 100 *µ*m. (**b**) Example direction-tuning curves of neurons located in putative L2/3 (*top*) or L6 (*bottom*), during V1 suppression (*blue*) and under control conditions (*black*). Drifting gratings with temporal and spatial frequencies coarsely optimized for the recording were presented for 0.75 s with continuous photostimulation starting 0.1 s before stimulus onset and lasting for 0.85 s. (**c**) Fold change (i.e. log2 ratio of average firing rates for V1 suppression and control conditions across tuning experiments with parameters as described in b)) as a function of cortical depth relative to the base of L4, estimated by CSD^57^. Neurons with fold change > 1 (i.e. more than doubling their rate) were considered putative PV neurons directly driven by photostimulation, and excluded from the computation of the layer-wise mean. *Gold*: layer-wise mean; *pink*: example neurons. Error bars: confidence intervals of the mean, determined by bootstrapping. n = 197 neurons. (**d**) Schematic size-tuning curve. (**e**) Recordings from dLGN. Responses of two example dLGN neurons to gray screen (size 0 deg) aligned to V1 suppression (*shaded blue*). *Red*: burst spikes, *black horizontal bar*: 200 ms. (**e**_1_): n = 54 trials; (**e**_2_): n = 105 trials. In all trials, speed ≤ 0:25 cm/s. (**f**) Firing rates during vs. before V1 suppression. n = 276 neurons; p = 1.5×10^-5^, Wilcoxon signed-rank test. (**g**) Ratio of burst spikes during vs. before V1 suppression. n = 232 neurons; p = 4:1×10^-11^, Wilcoxon signed-rank test. Data points at the margins represent neurons whose burst ratio was 0. *Inset*: cumulative distribution of burst lengths during (*blue*) vs. before (*black*) V1 suppression (*p* = 3.6 10*^−^*^13^, two-sample Kolmogorov-Smirnov test). (**h**) Size-tuning curves of an example dLGN neuron. The vertical bars at the bottom indicate preferred size; straight horizontal lines indicate responses to blank screen (size 0 deg). *Blue*: V1 suppression, *black*: control condition. Drifting gratings with orientation, temporal and spatial frequencies coarsely optimized for the recording were presented for 0.75 s, continuous photoactivation started 0.04 s before stimulus onset and lasted for 0.85 s. (**i**) Means of RoG fits for the dLGN population (*n* = 33; shaded areas: standard error or the mean (s.e.m.)). (**j-m**) Comparison of V1 suppression to control conditions for (**j**) responses to a small-sized stimulus (*p* = 0.0018), (**k**) responses to a large-sized stimulus (*p* = 0.03), (**l**) the preferred size (*p* = 0.00024), and (**m**) the corresponding suppression indices (SI) (*p* = 7.4 x 10*^−^*^5^, all Wilcoxon signed-rank test, n = 33 neurons). (**b, h**): error bars: s.e.m. (**f, g, j-m**): *pink*: example neuron, *gold*: population mean. In panel (g), markers of the two neurons indicated in pink almost completely overlap.

L6CT neurons are known to have low firing rates^49^, and to control firing along the V1 cortical column^34, 50^, likely via a translaminar inhibitory interneuron^51, 52^. Hence, to avoid concerns that direct L6 photoactivation might induce aberrant response patterns or recruit intracortical circuits in a widespread way, we suppressed the activity of L6CT neurons instead. To this end, we employed a strategy of reliable and powerful global V1 suppression^53^, by exploiting reversible optogenetic activation of PV+ interneurons, the major class of V1 inhibitory interneurons^54, 55^ (**Figure 2a**). We selectively targeted PV+ neurons for expression of ChR2 by injecting Cre-dependent AAV into V1 of PV-Cre mice. By recording extracellular activity across the layers of V1, we characterized V1 tuning for stimulus size and surround suppression (**Figure S4**), and, importantly, verified that optogenetic activation of PV+ neurons indeed suppressed output across the cortical column (**Figure 2b,c**). We found that - even in the presence of drifting gratings, which powerfully drive V1 activity under control conditions (**Figure 2b**) - optogenetic activation of PV+ inhibitory interneurons led to suppression of responses in V1 neurons across all layers (**Figure 2c**, 0 not included in 95% confidence intervals).

Having confirmed that photostimulation of PV+ interneurons suppressed V1 activity, even in infragranular layers, we turned to the thalamus and recorded from the dLGN (**Figure 2e-m**). Because dLGN RF locations in single electrode penetrations vary widely across simultaneously recorded neurons (e.g., **Figure 1h**), measuring complete size-tuning curves for dLGN neurons with RF-centered stimuli is laborious. We therefore decided to focus first on conditions without a visual stimulus, corresponding to 0 deg conditions in size-tuning experiments, and again restricted our analysis to trials without locomotion (speed *≤* 0.25 cm/s; **Figure 2d**). In light of a previous study showing that mouse dLGN responses to full-screen gratings during V1 suppression were enhanced^34^, we were surprised to observe that, in response to a uniformly gray screen (corresponding to a 0 deg size stimulus), suppression of the visual cortex for both shorter (250 ms; **Figure 2e**_1_) and longer (1 s; Figure 2**e**_2_) periods resulted in reduced geniculate activity. Indeed, firing rates of dLGN neurons during the window of V1 suppression were lower than in the same-sized window before V1 suppression (during: 3.7 sp/s vs. before: 4.9 sp/s; n = 276 neurons; *p* = 1.5 *×* 10*^−^*^5^, Wilcoxon signed-rank test; **Figure 2f**). Furthermore, closer inspection of the spike patterns (**Figure 2e**) centered around V1 suppression revealed a change in dLGN spiking: more specifically, the fraction of spikes fired in bursts (red) approximately doubled (during: 12.7% vs. before: 5.1%; n = 232 neurons; *p* = 4.1 *×* 10*^−^*^1^^1^, Wilcoxon signed-rank test; **Figure 2g**). In addition, V1 suppression shifted the entire distribution of burst lengths towards higher values (*p* = 3.6 *×* 10*^−^*^1^^3^, two-sample Kolmogorov-Smirnov test), including the median burst length (before V1 suppression: median 2 sp/burst, n = 835 bursts; during V1 suppression: median 3 sp/burst, n = 1739 bursts, *p* = 1.7 *×* 10*^−^*^1^^8^, Mann–Whitney U test; **Figure 2g, inset**). Both the decrease in firing rates and the subsequent increase in burst-spike ratio and burst length are consistent with the interpretation that, in the absence of stimulus drive, V1 suppression results in a decrease in feedback-mediated excitation. Such removal of excitation should hyperpolarize dLGN cells, resulting initially in fewer action potentials and later (*≥* 100 ms) in bursting, given the hyperpolarization-mediated de-inactivation of T-type calcium channels. T-type calcium channels, which are abundant in the thalamus, mediate low-threshold calcium spikes, whose amplitude is inversely related to membrane potential and is correlated with the number of action potentials in a burst riding its crest^56^. Complementary to the results of global V1 suppression, we found that photoactivation of L6CT neurons under size 0 deg conditions (i.e., absence of sensory stimulation) was sufficient to enhance tonic firing in dLGN (**Figure S5**). Overall, in the absence of visual stimuli, CT feedback seems to exert its effect mainly through the direct, excitatory pathway, boosting firing rates and promoting the tonic firing mode.

We next presented drifting gratings of various sizes centered on the RF of each dLGN neuron and recorded responses while interleaving trials with and without optogenetic suppression of V1 (**Figure 2h-m**). To obtain size-tuning curves, we fit each dLGN neuron’s responses to gratings of various sizes, under either the control or V1 suppression condition, with a descriptive model of size tuning (ratio-of-Gaussians model, RoG)^58^ that defines the relative contributions of an excitatory center and inhibitory surround (**Figure 2d, h-i**, **Figure S6**, only trials without locomotion). From this fit, we determined the preferred size as the size that elicited the maximum response, and the suppression strength as an index (SI) that quantifies the response to the surround relative to the center, with 0 indicating no suppression and 1 indicating full suppression.

In order to probe the effects of cortical feedback on spatial integration, we first asked if V1 suppression differentially affected dLGN spiking responses to drifting gratings according to stimulus size. For small stimuli (i.e., the modelled response at the preferred size under the control condition), we observed, as in the case of spontaneous activity, that V1 suppression caused a decrease in dLGN responses (control: 22.34 sp/s vs. V1 suppression: 18.09 sp/s, n = 33 neurons; *p* = 0.0018, Wilcoxon signed-rank test; **Figure 2j**). However, for large sizes (i.e., the modelled response to a 200 deg stimulus), we found, in accordance with previous results^34^, the opposite effect: here, dLGN responses increased during V1 suppression (control: 11.69 sp/s vs. V1 suppression: 13.22 sp/s, n = 33; *p* = 0.033, Wilcoxon signed-rank test; **Figure 2k**). Hence, our results indicate that cortical feedback can affect dLGN responses in a contextual manner, enhancing responses to the preferred size while suppressing responses to larger stimuli. To determine whether the size-dependent modulation of dLGN firing rates during V1 suppression translated into significant changes in spatial integration, we next examined the size-tuning curves of individual cells. Indeed, we found that during V1 suppression, dLGN neurons preferred larger sizes (control: 16.27 deg vs. V1 suppression: 20.42 deg; n = 33; *p* = 0.00024, Wilcoxon signed-rank test; **Figure 2l**) and were less surround-suppressed (control: 0.62 vs. V1 suppression: 0.45; n = 33; *p* = 7.36 *×* 10*^−^*^5^, Wilcoxon signed-rank test; **Figure 2m**).

While V1 suppression did not abolish surround suppression in the dLGN (28/33 cells still had *SI ≥* 0.1), our results indicate a substantial involvement of cortical feedback in shaping spatial integration in dLGN: feedback enhances contextual effects, facilitating responses to the center while suppressing those to the surround, which results in sharper RFs and a stronger center-surround antagonism. While the enhanced small-size responses are consistent with a net depolarizing effect of CT feedback, the increased surround suppression for large sizes is suggestive of CT feedback acting via inhibition.

### Capturing effects of feedback on dLGN spatial integration requires widespread CT inhibition in a firing rate model

How might CT feedback shape dLGN spatial integration via inhibition? We first investigated this question with the aid of a previously developed mechanistic firing-rate model of dLGN, the extended difference-of-Gaussians model (eDoG)^59, 60^. In the eDOG model (**Figure 3a**), the response of a dLGN relay cell (*R*) arises from excitatory and inhibitory feedforward input from retinal ganglion cells (*G*) whose RF is described by a difference-of-Gaussians model^61^, and excitatory and inhibitory feedback input from a population of cortical neurons (*C*) with various orientation preferences. The connections between the three processing stages are represented by 2D Gaussian spatial coupling kernels (*K*), whose amplitude captures the synaptic strength, and whose width captures the spatial scale over which visual information is integrated. The model is segregated into an ON and OFF pathway, where feedforward input is pathway specific, while cortical feedback is pathway unspecific (cross-symmetric), arising from both ON and OFF pathways. While the eDOG model makes clear simplifications, for instance by not explicitly considering intracortical connectivity or transformations^59^, its conceptually and mathematically simple framework and the availability of a simulation tool^60^ make it a prime choice. We adjusted the model’s parameters to reflect known properties of the mouse visual system (**Table S1**). While the model is agnostic to the source of feedback-mediated inhibition, it permits exploration of how the spatial scale of inhibitory feedback shapes dLGN spatial integration.

**Figure 3.**
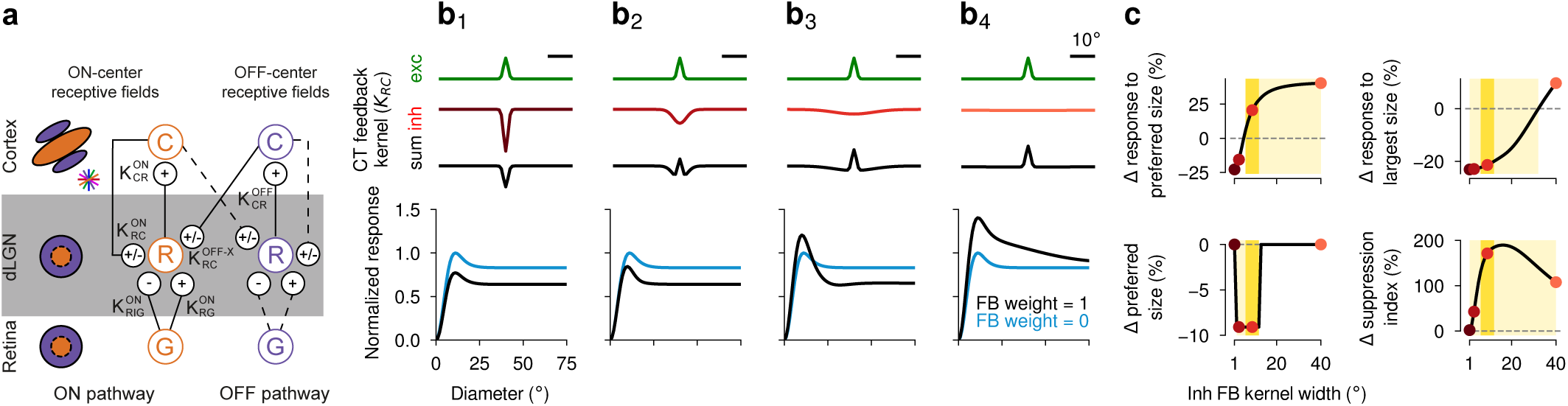
A wide inhibitory feedback coupling kernel is necessary to reproduce feedback-enhanced surround suppres- sion and sharpening of RFs observed in the dLGN. (**a**) Schematic of the extended difference-of-Gaussians model (eDOG). In the eDOG model, center-surround RFs of dLGN relay cells (*R*) are modelled by feedforward inputs from retinal ganglion cells (RGCs, *G*) and feedback inputs from cortex (*C*). The three layers are connected by 2D Gaussian coupling kernels (*K*). RGCs, whose RFs are described by a difference-of-Gaussians model, provide excitatory (*K_RG_*) and inhibitory (*K_RIG_*) feedforward input to relay cells. CT feedback of both signs (*K_RC_*) originates from a population of cortical cells, which in turn receive input from dLGN relay cells (*K_CR_*). While the feedforward pathway is segregated into ON and OFF pathways, CT feedback is cross-symmetric, meaning that e.g. ON center relay cells receive excitatory and inhibitory feedback from cortical ON and OFF cells. Although cortical cells are orientation tuned, a given relay cell receives input from a diverse population of cortical cells (colored bars), making the net effect of CT feedback insensitive to the orientation of the stimulus. Adapted from Mobarhan et al. (2018)^60^. (**b**) Effects of varying the width of the inhibitory CT feedback coupling kernel on simulated size-tuning curves. *Top*: excitatory (*green*) and inhibitory (*red*) CT feedback coupling kernels, and their sum (*black*). (**b**_1_) Same width (*σ*_inh fb_ = *σ*_exc fb_), (**b**_2_) same width as inhibitory feedforward kernel (*σ*_inh fb_ = *σ*_inh ff_), (**b**_3_**, b**_4_) two larger widths (**b**_3_: *σ*_inh fb_ = 9 × *σ*_exc fb_; **b**_4_: *σ*_inh fb_ = 40 × *σ*_exc fb_). Note that the area under the curve is the same for all inhibitory feedback coupling kernels. *Bottom*: simulated dLGN size-tuning curves with cortical feedback intact (*black*) and abolished (*blue*). For each width, responses are normalized to the peak response seen under the condition without CT feedback. (**c**) Effects of CT feedback for kernel widths between 1 deg and 40 deg (varied in 1 deg steps) on the response magnitude to the preferred stimulus size (*top left*), the response magnitude to the largest stimulus (*top right*), the preferred size (*bottom left*), and surround suppression (*bottom right*), revealed by zeroing the weight of CT feedback. *Red* points correspond to kernel widths depicted in **b**. The parameter ranges within which simulations yield results that are qualitatively similar to experimental observations are indicated with regard to each single variable (*light yellow*) vs. all four variables (*dark yellow*). Light yellow ranges correspond to 6 deg *≤ σ*_inh fb_ *≤* 40 deg (response to preferred size), 1 deg *≤ σ*_inh fb_ *≤* 32 deg (response to largest size), 2 deg *≤ σ*_inh fb_ *≤* 12 deg (preferred size), 1 deg *≤ σ*_inh fb_ *≤* 40 deg (suppression index). The dark yellow range corresponds to 6 deg *≤ σ*_inh fb_ *≤* 12 deg.

To assess how inhibition via CT feedback might increase surround suppression and sharpen the RFs of dLGN neurons, we systematically varied the width of the inhibitory feedback coupling kernel (**Figure 3b, top**) and simulated tuning curves for grating patches of different sizes, with and without CT feedback (**Figure 3b, bottom**). When we set the width parameter of the inhibitory CT feedback kernel to equal the width of the excitatory CT feedback kernel (*σ*_inh fb_ = *σ*_exc fb_), the model failed to replicate our experimentally observed results (**Figure 3b**_1_): CT feedback had an overall suppressive effect, reducing responses for all stimulus sizes (22.8% decrease for preferred stimulus size; 23.1% decrease for largest stimulus size) and failed to substantially alter the preferred stimulus size and surround suppression (1.9% increase). We then increased the spatial scale of inhibitory CT feedback to match the spatial scale of feedforward inhibition (*σ*_inh fb_ = *σ*_inh ff_, **Figure 3b**_2_). Now CT feedback began to decrease the preferred size (9.1% decrease) and increase surround suppression (42.7% increase), but it still led to weaker responses overall, even for small sizes (15.6% decrease for preferred stimulus size; 22.9% decrease for largest stimulus size). Only when the width of the inhibitory CT feedback component was sufficiently large (*σ*_inh fb_ = 9 *× σ*_exc fb_; **Figure 3b**_3_) did our simulations yield a pattern comparable to the size-dependent effects observed on average in the experimental data: while responses to the preferred stimulus size were enhanced (20.6% increase), responses to the largest stimulus size were suppressed (21.4% decrease). In accordance with the experimental data, we also observed that CT feedback decreased the preferred size (9.1% decrease) and strengthened surround suppression (171.2% increase). Finally, when we further increased the spatial scale of the inhibitory feedback kernel (*σ*_inh fb_ = 40 *× σ*_exc fb_; **Figure 3b**_4_), CT feedback increased firing rates independently of stimulus size (40.4% increase for preferred stimulus size; 9.7% increase for largest stimulus size) and enhanced surround suppression (107.7% increase), but did not affect the preferred stimulus size (0.0% change).

Analysis of simulations with more extensive variation of the width of the inhibitory CT feedback kernel revealed that feedback-induced amplification of responses to the preferred size (**Figure 3c**, top left, light yellow) and strengthening of surround suppression (**Figure 3c**, lower right, light yellow) required sufficiently wide kernels. Much wider kernels, however, failed to reproduce both the feedback-induced decrease of responses to larger stimulus sizes (**Figure 3c**, top right, light yellow) and the sharpening of RFs (**Figure 3c**, lower left, light yellow), which restricted the parameter range that replicated our average experimental results to larger but spatially confined inhibitory feedback kernel widths (**Figure 3c**, dark yellow). Taken together, the model suggests that cortical feedback enhances contextual effects in dLGN via an inhibitory component that integrates information over a sufficiently large, yet still localized, spatial scale.

### RF properties of mouse visual TRN are suited for providing wide-scale inhibition to dLGN

A candidate circuit through which the cortex could exert widespread inhibitory influence over the dLGN is indirect inhibition via the visual TRN (visTRN, **Figure 4**). Present in all mammals, the TRN comprises a sheath of GABAergic neurons surrounding the lateral and anterior parts of the thalamus^62, 63^. Since TRN receives input from axon collaterals of both thalamic relay cells and CT neurons, it is in a prime position to modulate the flow of information between the thalamus and the cortex^62, 63^. Owing to its inhibitory projections to dLGN, the visTRN has been considered a “guardian of the gate to the cortex”^64^, and has been implicated in gain control^64, 65^ and attentional selection^66–70^.

**Figure 4.**
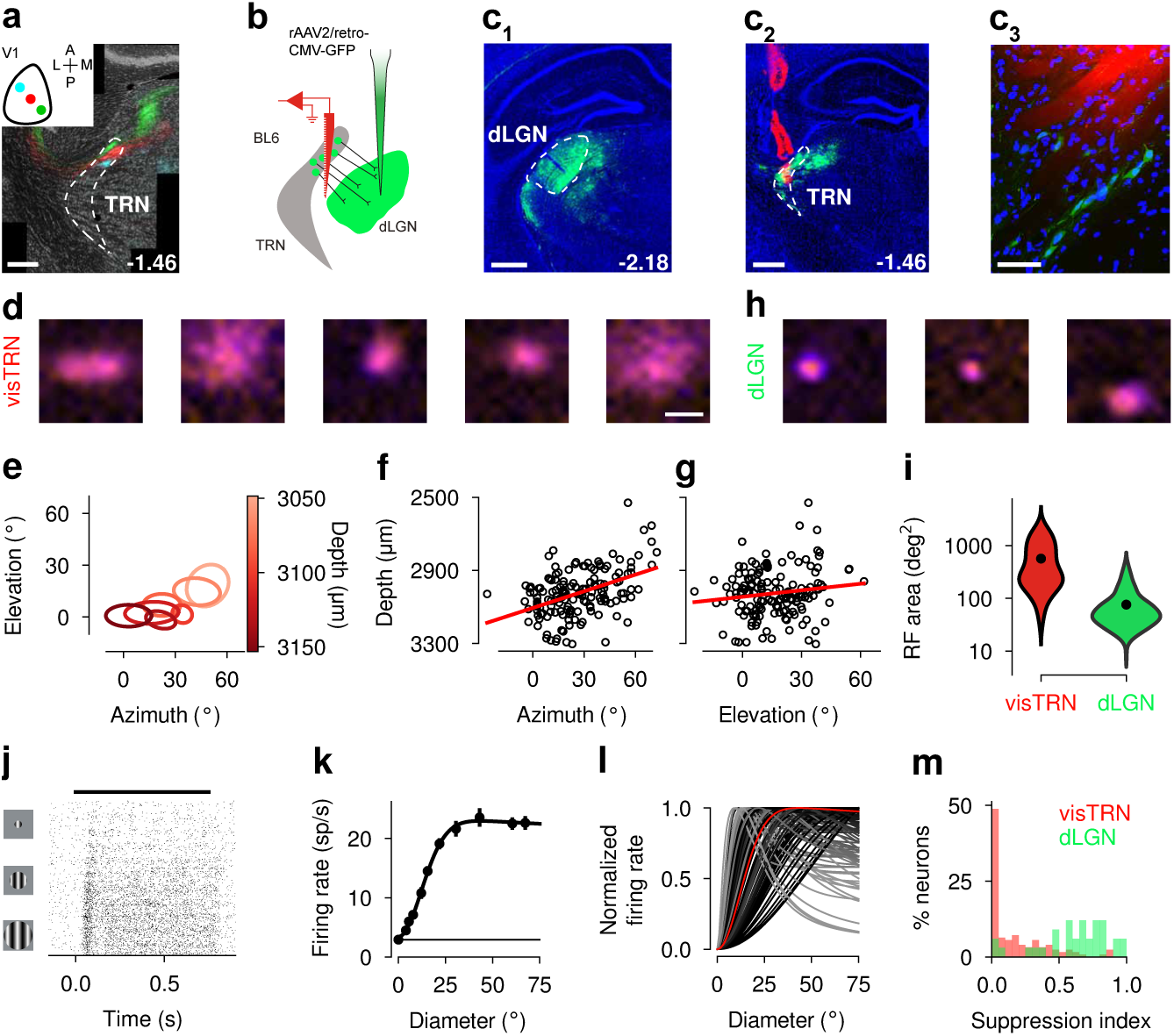
Retinotopic organization and RF properties of visTRN neurons. (**a**) V1 L6CT terminal fields in visTRN, obtained from the triple-color viral injections along the V1 azimuth axis (see also Figure 1a, **Figure S7**). dashed outline: TRN; scale bar: 0.25 mm; numbers indicate distance from bregma in mm. Inset: cartoon of V1 injection pattern. (**b**) Schematic of viral labeling of visTRN neurons with GFP by injecting a retrograde AAV^72^ into the dLGN. (**c**) GFP labeling of visTRN neurons in an example mouse. (**c**_1_): dLGN injection site. Dashed outline: dLGN; scale bar: 0.5 mm. (**c**_2_): visTRN neurons labeled by retrograde AAV and visTRN recording site. Dashed outline: TRN; scale bar: 0.5 mm. (**c**_3_): Magnified view around the tip of the electrode trace from the slice shown in (c_1_). Scale bar: 50 *µ*m. All panels: *blue*: DAPI, *green*: GFP, *red*: DiI-labeled electrode trace. (**d**) Classical RFs for five example visTRN neurons. *Orange*: OFF response, *purple*: ON response, scale bar: 20 deg. (**e**) RF fits from an example visTRN recording session (n = 7 simultaneously recorded neurons). (**f**) Comparison of the recording depth and the azimuth coordinate of the RF center for the visTRN population (n = 154). *Red line*: linear regression, *R*^2^ = 0.2; *p* = 5.43 x 10*^−^*^9^ (slope). (**g**) Same as (f) but for the elevation coordinate of the RF center. *R*^2^ = 0.02; *p* = 0.08. (**h**) Classical RFs for three dLGN example neurons. (**i**) Comparison of classical RF sizes for recorded visTRN (n = 218) and dLGN (n = 197) neurons. Outlines indicate distribution of classical RF sizes. *Black*: mean; *p* = 1.0 10*^−^*^51^, Mann-Whitney U test. (**j**) Raster plot of an example visTRN neuron recorded in a size-tuning experiment. Trials are sorted by stimulus size with lower rows showing responses to larger stimulus sizes. 50 trials per size; black horizontal bar: stimulus presentation period. (**k**) Size-tuning curve corresponding to (j). Horizontal line: response to size 0 deg. Error bars: s.e.m.. (**l**) Size-tuning curves for visTRN cell population (n = 125). Strength of surround suppression is represented by darkness of line. *Red*: example neuron from (j,k). (**m**) Distribution of suppression indices for the recorded visTRN population (*red*) and the dLGN population (*green*).

To better understand how feedback signals from primary visual cortex arrive in visTRN, we first characterized the organization of V1 L6CT inputs by analyzing visTRN slices obtained in our triple-color viral tracing experiment (**Figure 4a**). For injections along the V1 azimuth axis, we found clearly separated, topographically organized terminal fields within single coronal slices (**Figure 4a**). For injections along the V1 elevation axis, the differently colored terminal fields in visTRN were distributed along the anterior-posterior (A-P) axis (**Figure S7**).

To explore whether CT feedback might enhance surround suppression in the dLGN via the visTRN, we next tested whether mouse visTRN neurons have appropriate feature selectivity, i.e. large, retinotopically organized RFs, responses that increase with stimulus size, and little surround suppression. We recorded from visTRN by inserting a silicon probe (positioned at an appropriate stereotaxic location^71^) to a depth of *∼* 3500 *µ*m (**Figure 4b**) until we found neurons with vigorous, visually evoked responses. Since visTRN is located near other thalamic nuclei with visually responsive neurons, we confirmed *post-mortem* via retrograde viral labeling that our visTRN recording sites were in the vicinity of neurons that provide input to dLGN (**Figure 4c**). Indeed, after injection of rAAV2/retro-CMV-GFP^72^ into the dLGN (**Figure 4c**_1_), we found dense GFP expression in the dorsocaudal part, corresponding to the visual sector of TRN^62, 70, 73^ (**Figure 4c**_2_). Closer inspection revealed retrogradely labeled cell bodies, localized near the DiI-labeled electrode track (**Figure 4c**_3_). This histological evidence, in combination with the robust visual responses encountered during our recordings, confirmed that we had indeed targeted the visTRN.

To test the RF properties of mouse visTRN, we first mapped classical RFs of single visTRN neurons using a sparse noise stimulus (**Figure 4d, left**). RFs of visTRN neurons covered a wide range of sizes, with individual neurons displaying small (**Figure 4d**, second from the right: area = 169.3 deg^2^; *R*^2^ = 0.92) or large RFs (**Figure 4d**, rightmost: area = 780.5 deg^2^; *R*^2^ = 0.84). Focusing on RFs obtained within a single penetration (**Figure 4e**), we realized that fitted RF centers followed a coarse retinotopy, with neurons recorded at deeper, more ventral electrode sites having more central RF locations (**Figure 4e**). Both RF azimuth (n = 154; *p* = 5.43 *×* 10*^−^*^9^; **Figure 4f**) and elevation (*p* = 0.08; **Figure 4g**) predicted recording depth, but the relationship was considerably stronger for azimuth (*p* = 0.028, ANCOVA). Importantly, the retinotopic organization of visTRN neurons seems to match the topographic arrangement of the V1 L6CT feedback terminal fields (compare to **Figure 4a**, **Figure S7**). Injections along the V1 azimuth produced L6CT axonal labels within a coronal plane in visTRN, with central visual space being represented most ventrally, while injections along the V1 elevation axis produced labels predominantly along the A-P axis, which we cannot capture with individual single-shank recordings.

While the overall match between retinotopy of the CT innervation and visTRN organization is consistent with preserving spatial information, our simulations predicted inhibitory feedback to be spatially extensive, so we next focused on visTRN RF size. Comparing the classical RF size of visTRN (n = 218 neurons; 566.1 *±* 37.4 deg^2^; mean *±* s.e.m.; examples in **Figure 4d**) with a population of dLGN neurons measured under the same conditions (n = 197; 75.9 *±* 5.1 deg^2^; examples in **Figure 4h**) revealed that, despite having overlapping distributions, classical RFs of visTRN neurons were on average 7.5*×* larger (*p* = 1.0 *×* 10*^−^*^5^^1^, Mann-Whitney U test) and more variable in size (*p* = 8.3 *×* 10*^−^*^2^^3^, Brown-Forsythe test, **Figure 4i**).

Finally, centered on the RFs, we presented drifting gratings of various sizes and fit the averaged responses obtained in trials without locomotion (run speed < 0.25 cm/s for more than half of the trial duration) with the RoG model (**Figure 4j-m**). Analogous to our analysis of dLGN size tuning, we used the model fit to determine, for each visTRN neuron, its preferred size and the strength of surround suppression. Like the example shown (SI: 0.02, **Figure 4j, k**), the majority of visTRN neurons experienced little to no surround suppression (n = 125 visTRN neurons; mean SI: 0.17 *±* 0.02, **Figure 4l, m**). In fact, almost half of the population (48.8%) had an SI less than 0.05. In comparison to dLGN neurons (n = 33 dLGN neurons; mean SI: 0.62 *±* 0.05), visTRN neurons experienced significantly less surround suppression (*p* = 2.8 *×* 10*^−^*^1^^2^, Mann-Whitney U test, **Figure 4m**). Thus, similar to neurons in visTRN of carnivores and primates (perigeniculate nucleus)^74–78^, mouse visTRN neurons follow a coarse retinotopic organization, have spatially localized yet large RFs, and experience little surround suppression themselves. By responding weakly to small stimuli and strongly during presentation of large stimuli, the properties of inhibitory visTRN neurons are well suited for sculpting dLGN surround suppression.

### Suppressing cortical feedback increases preferred size and reduces responses in visTRN, in particular for large stimuli

If CT feedback indeed enhances dLGN surround suppression by recruiting inhibition from visTRN, how might CT feedback influence visTRN responses? Our model, where scaling the amplitude of a spatially fixed set of excitatory and inhibitory feedback coupling kernels is sufficient to reproduce all experimentally observed changes of dLGN size tuning, makes one specific prediction: If CT feedback provides substantial indirect inhibition via the visTRN, then suppression of V1 should reduce visTRN responses.

To test this hypothesis, we measured the responses of visTRN neurons in PV-Cre mice to drifting gratings of varying size, with interleaved trials in which we suppressed CT feedback by photoactivating PV+ inhibitory interneurons in V1 (**Figure 5a**). When we inspected the raster plots (**Figure 5b**) and fitted size-tuning curves (RoG model) of single visTRN neurons (**Figure 5b**, **Figure S8**, again focusing on trials without locomotion), we found that suppressing V1 reduced overall responsiveness. This reduction was robust not only for the neuron shown in **Figure 5b,c** (-46.2%), but also for the population of recorded visTRN neurons (V1 suppression: 15.6 *±* 2.2 sp/s; control: 23.4 *±* 2.5 sp/s; n = 67; *p* = 4.9 *×* 10*^−^*^10^; Wilcoxon signed-rank test; **Figure 5d,e**). Similar to our findings in dLGN, suppressing V1 also increased visTRN neurons’ burst ratios (V1 suppression: 12.8%; control: 10.1%; n = 67 neurons; *p* = 0.006; Wilcoxon signed-rank test; **Figure 5f**). We conclude from the substantial reduction in responsiveness during V1 suppression that visTRN is strongly engaged by CT feedback.

**Figure 5.**
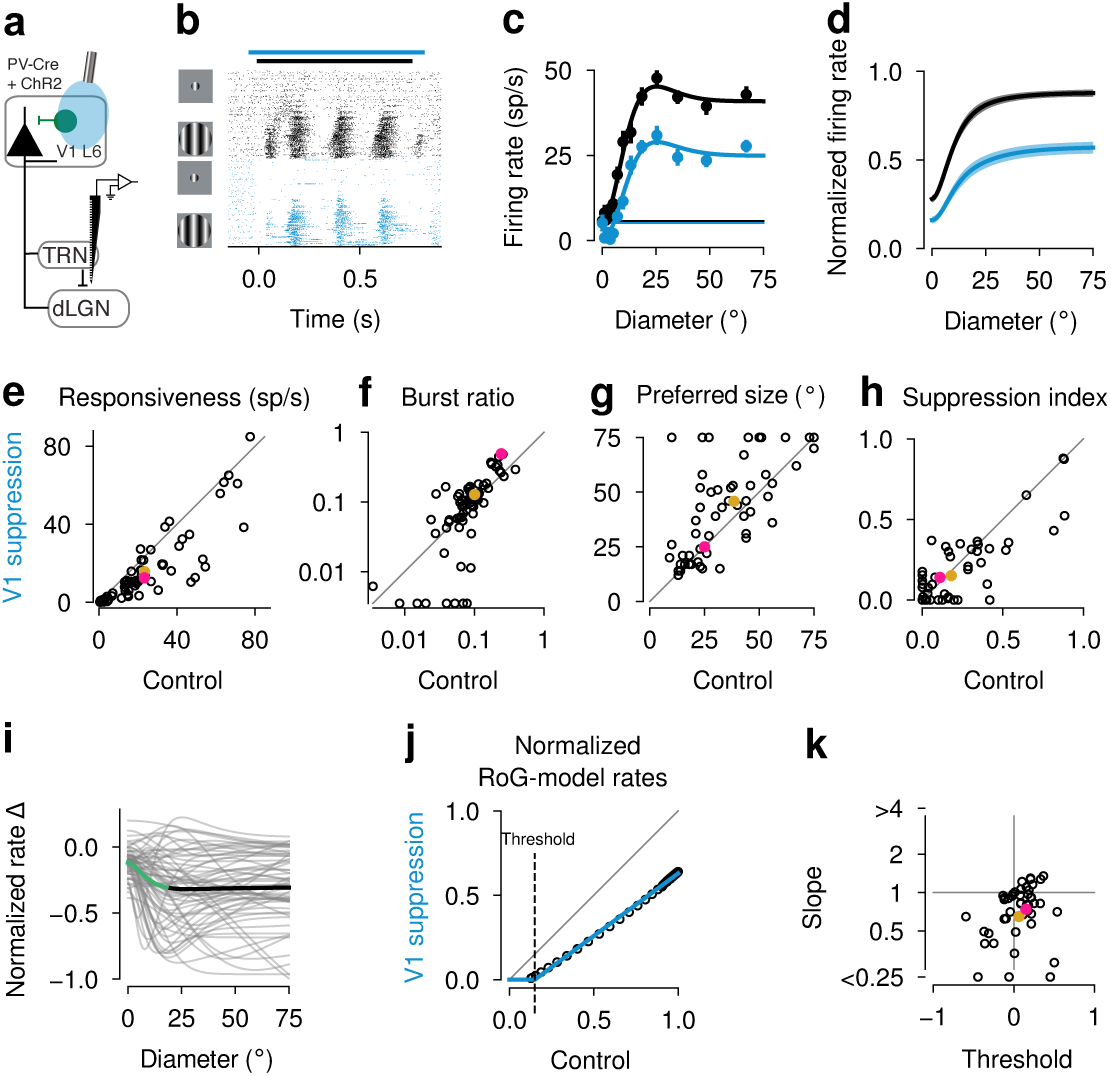
Suppression of cortical feedback reduces responses and increases preferred stimulus size in the visTRN. (a) Schematic of the experimental approach. (**b**) Raster plot of a visTRN neuron recorded in a size-tuning experiment. Trials are sorted by feedback condition and stimulus size, with lower rows depicting responses to larger stimulus sizes (20 trials per size and feedback condition; black horizontal bar: stimulus presentation period; blue horizontal bar: continuous V1 suppression period). Drifting gratings with temporal and spatial frequencies coarsely optimized for the recording were presented for 0.75 s with continuous photostimulation starting 0.04 s before stimulus onset and lasting for 0.85 s. (**c**) Size-tuning curves, same neuron as in **b** (horizontal bars: response to size 0 deg; error bars: s.e.m.). (**d**) Means of RoG fits for the visTRN population (*n* = 63; shaded areas: s.e.m.). (**e-h**) Mean evoked response (**e**; n = 67; *p* = 4.9 x 10*^−^*^10^), burst ratio (**f**; n = 67; *p* = 0.006), preferred size (**g**; n = 61; *p* = 0.001) and suppression indices (**h**; n = 61; *p* = 0.18; Wilcoxon signed-rank test) for visTRN cell population. In (f) data points at the margins represent neurons whose burst ratio was 0. (**i**) Difference between normalized ratio-of-Gaussians models for V1 suppression and control conditions (*gray*: single visTRN neurons; *black*: population mean; *green*: size range, in which a 1 deg increase in size led to a significant increase in the CT feedback effect (0 deg - 18 deg; *p <* 0.05; bootstrap test; *n* = 63); for sizes *>* 18 deg, the CT feedback effect plateaus. (**j**) Threshold linear fit (*blue*) to RoG model evaluated in 1-deg steps (*black*) for the visTRN neuron in (b, c) (slope: 0.74; threshold: 0.15; *R*^2^ = 1). (**k**) Slope (*p* = 2.8 x 10*^−^*^5^) and threshold (n = 46; *p* = 0.04, Wilcoxon signed-rank test) parameters extracted from threshold linear fits for the visTRN population. (**b-d**): *blue*: V1 suppression; *black*: control; (**e-j, j**): *pink*: example neuron; *gold*: population mean.

More closely inspecting the parameters of the fitted size-tuning curves, we realized that the reduction of visTRN responses during V1 suppression was not uniform across stimulus sizes. Focusing on those visTRN neurons which were still responsive during V1 suppression (mean firing rate *≥* 0.1 sp/s, 61 of 67 neurons), we found that V1 suppression increased visTRN preferred size (V1 suppression: 45.8 *±* 2.9 deg; control: 38.5 *±* 2.7 deg; n = 61 neurons; *p* = 0.001; Wilcoxon signed-rank test; **Figure 5g**). This increase indicates that visTRN’s peak inhibitory output to dLGN might shift towards larger stimulus sizes, which could contribute, besides the overall reduction in strength of the inhibitory CT feedback component, to our observation that V1 suppression increased dLGN preferred size (**Figure 2l**). While we found the effects of CT feedback on visTRN to be overall consistent, the remaining variability across the visTRN population seems unrelated to several visTRN response properties (**Figure S9**).

While CT feedback did not change the overall strength of surround suppression in visTRN (SI during V1 suppression: 0.15 *±* 0.03; control: 0.18 *±* 0.03; n = 61; *p* = 0.18; Wilcoxon signed-rank test; **Figure 5h**), we found that modulation of visTRN responses by CT feedback nevertheless was not constant across stimulus sizes. Inspection of the differences in the normalized fitted size-tuning curves between the two conditions showed that responses to larger stimuli (*>* 18 deg) were more strongly affected by V1 suppression than responses to smaller stimuli (0–18 deg), where the effect of CT feedback steadily increased (n = 63; *p <* 0.05; bootstrap test; **Figure 5i**, green line; see also **Figure 5d**). Hence, while CT feedback seems to enhance visTRN responses across all stimulus sizes, this enhancement became progressively stronger with increasing stimulus size before reaching a plateau.

What is the nature of the transformation exerted by CT feedback on visTRN responses? If V1 suppression reduced visTRN responses regardless of stimulus size - and thus independently of the visTRN activity level - the effect would be best explained by a subtractive mechanism. However, since V1 suppression was more effective for large stimuli, which under control conditions evoked the strongest response, the effect might instead be based on a divisive mechanism. Indeed, the dynamic regulation of the amount of background barrages of CT synaptic input (“synaptic noise”)^79, 80^ has been proposed as a top-down mechanism for gain modulations in the thalamo-cortical system^81^.

To identify the computation governing visTRN responses during modulations of CT feedback, we fit a threshold-linear model (**Figure 5j**, blue), which predicts responses during V1 suppression by shifting and scaling responses observed under the control condition. Because V1 suppression cannot lead to negative firing rates, the model additionally zeroed firing rates up to a threshold for activation. Although it is impossible for this simple model to capture the observed changes in preferred size, it did capture the effects of V1 suppression on size-tuning curves reasonably well for the majority of visTRN neurons (46/63 neurons, *R*^2^ *≥* 0.8). Focusing on this subset of well-fit neurons, in which V1 suppression had mainly linear effects, we found for both the neuron shown in **Figure 5j** (*R*^2^ = 1; threshold: 0.15; slope: 0.74; same neuron as in **Figure 5b,c**) and the recorded population as a whole (**Figure 5k**) a mild subtractive effect (threshold: 0.06 *±* 0.04, mean *±* s.e.m.; *p* = 0.04, Wilcoxon signed-rank test) and a substantial and consistent divisive effect (slope: 0.65 *±* 1.13; *p* = 2.8 *×* 10*^−^*^5^, Wilcoxon signed-rank test; **Figure 5k**). Since divisive scaling implies that high firing rates are reduced most, and visTRN neurons have high responses to large stimuli (see **Figure 4j-m**), this analysis further corroborates our finding that CT feedback most strongly engages visTRN activity in response to large stimuli. Such size-dependent recruitment of inhibition via the visTRN by CT feedback might account for our earlier finding that the dLGN’s responses to large stimuli are enhanced when CT feedback is suppressed (**Figure 2k**). Taken together, the substantial modulation of visTRN responses, and the size-dependent recruitment of inhibitory visTRN responses by CT feedback, make visTRN an ideal candidate for mediating feedback-enhanced surround suppression in dLGN.

## Discussion

Using a combination of viral tracing, bidirectional optogenetic manipulations, and computational modeling, we have shown that one role of the retinotopically organized cortical feedback to the mouse dLGN is to sculpt spatial integration by sharpening RFs and enhancing surround suppression. We identified spatially specific, distant suppressive influences of corticothalamic feedback, which are most consistent with indirect inhibition. In accordance with simulations in our thalamocortical network model, which indicated that widespread inhibitory CT feedback is required to reproduce our experimental results, we show that the spatial selectivity of neurons in visTRN and their size-specific recruitment by CT feedback make them an ideal candidate for mediating feedback-enhanced surround suppression in dLGN. Therefore, corticothalamic feedback, most probably with the involvement of the TRN, sharpens spatial responses and strengthens contextual modulations in dLGN.

### Spatial integration in dLGN

Spatial integration in the dLGN is achieved by multiple mechanisms, as surround suppression occurs both up- and downstream of dLGN. Indeed, it is first established in the retina^82–84^, and is also a hallmark of responses in area V1^43, 44, 85–89^. The mechanisms underlying surround suppression in dLGN therefore include inheritance from feedforward retinal input^26, 90^, augmentation via non-linearities at the retino-geniculate relay^91^, recurrent thalamic inhibition^92, 93^, and CT feedback^28, 29, 31, 33, 94^. The CT feedback-mediated sharpening of RFs and strengthening of the center-surround antagonism that we found in dLGN of awake mice parallels earlier results in anesthetized cats^28, 30, 31^, ferrets^32^, and non-human primates^27, 29, 33^.

While the underlying circuit mechanisms might differ, the effects of corticothalamic feedback on spatial integration have similar signatures to those of corticocortical feedback. Indeed, inactivating higher visual cortical areas in both mouse and primate cortex reduces surround suppression in V1 neurons^95–98^ and shifts their preferred sizes towards larger stimuli^95, 96^. This suggests that the sculpting of spatial integration could represent a canonical function of feedback in the visual system.

Furthermore, corticothalamic feedback has similar effects on sharpening stimulus selectivity in other sensory modali- ties^11, 99–103^. In the rat somatosensory system, for instance, pharmacological activation of layer 6 neurons increases responses in topographically aligned thalamic barreloid neurons, while it reduces responses in adjacent, topographically misaligned barreloid neurons^99^. Similarly, in the bat echolocation system, corticofugal projections sharpen and adjust auditory frequency tuning^101^. This indicates that the effect of corticothalamic feedback on spatial integration generalizes across different sensory modalities.

### The role of the TRN

By measuring RF properties in the visTRN and their modulation by CT feedback, and by simulating the impact of inhibitory feedback at various spatial scales in a mechanistic dLGN model^60^, we found evidence that CT feedback can sculpt dLGN spatial integration via visTRN. While visTRN has long been implicated in controlling the dLGN^64^, the specific form of this control has been a matter of debate, ranging from homogenizing dLGN activity (“thermostat hypothesis”) to triggering focal re-bound excitation in dLGN (“searchlight hypothesis”)^64, 104^. Although our results share with the “searchlight hypothesis” a component of spatial specificity, they are different by implying that dLGN spatial selectivity might be enhanced by direct, localized excitation from L6CT pyramidal cells acting in concert with indirect, more widespread inhibition from visTRN.

An alternative source of indirect inhibition in dLGN are local interneurons, which make up *∼* 5% of mouse dLGN^105^ and whose connectome allows them to participate in both local and global inhibitory processing^106^. Studies in cats have shown that dLGN interneurons are sensitive to polarity, are organized into concentric subunits of opposite sign^107^ and provide dLGN relay cells with either specific opposite-sign (“push-pull”) or same-sign inhibition^107, 108^, all of which argues in favor of local influences. However, since signaling in dLGN interneurons can occur via dendrites without depolarization of the cell body^109^, the RF properties of dLGN interneurons measured as spiking output likely represent only a subset of the filtering operations that this neuron type can perform. To disentangle the relative contributions of dLGN interneurons and the visTRN to feedback-enhanced surround suppression in dLGN, targeted recordings from geniculate interneurons, and an assessment of their modulation by CT feedback will be a crucial next step.

Our finding that V1 suppression modulates firing rate in the dLGN, while substantially reducing visTRN responses, sheds further light on the role of cortical vs. subcortical inputs for these two thalamic nuclei. Our estimation of a *∼* 50% V1 contribution to visTRN firing rates during size tuning fits observations in slice preparations, where EPSPs elicited by stimulation of L6CT neurons are larger in the TRN than in relay neurons^110, 111^. The impact of CT pyramidal cell input to visTRN *in vivo*, however, is less clear. Similar to our results, some previous studies noted a strong reduction of visTRN activity during CT feedback suppression^36, 112, 113^; others, however, observed no changes in visTRN responses upon removal of CT feedback, and therefore concluded that visTRN was mainly driven by subcortical inputs^78, 114^. A possible explanation for this discrepancy might be that in anesthetized animals the effects of CT feedback on visTRN responses have been underestimated, because the responsiveness of feedback projections^115^, including CT feedback^18^, might be particularly reduced during anesthesia, and attentional processes adding to the normal recruitment of CT feedback are lacking^70, 113^.

Besides the pronounced reduction in responsiveness of visTRN neurons during V1 suppression, we also observed an increase in preferred size. While we do not have any direct evidence to explain this finding, our data is consistent with at least two mechanisms: increased visTRN preferred size during V1 suppression might simply reflect the increased preferred size of dLGN providing feed-forward input to visTRN. In addition, since V1 suppression promotes bursting also in visTRN, electrotonic coupling of the TRN network through electrical synapses^62, 116^ might be more prevalent^117^: electrical synapses would allow propagating the slow low-threshold spike component of the burst more efficiently than fast sodium action potentials^118^, which might increase the spatial spread of activity within the visTRN network.

Although TRN exclusively consists of GABAergic neurons, their anatomical and functional composition is diverse. Recent evidence suggests that, similar to primates and cats^119, 120^, the sensory sections of mouse TRN can be further divided into lower and higher-order subnetworks^73, 121, 122^. For instance, SOM+ neurons located at the edges of mouse TRN have been found to preferentially innervate higher-order thalamic nuclei. These neurons receive little input from primary sensory cortices and are less likely to fire bursts of action potentials. In contrast, PV+ neurons located in the central portion of TRN project to first-order thalamic nuclei, receive input from primary sensory cortices, and are more likely to fire in long bursts^73, 121^. While cortical SOM+ neurons have been implicated in strengthening surround suppression in V1^89^, SOM+ TRN neurons might thus not play an important role for CT feedback mediated surround suppression in dLGN. Furthermore, the influence of indirect CT feedback via visTRN onto dLGN is likely behavioral state-dependent^68^. While we focused our analyses on the quiescent state, it will be interesting for future studies to further explore dynamic, state-dependent changes in the activity and selectivity of V1 L6CT^123^ and visTRN neurons^68^.

Our findings regarding the role of CT feedback in shaping dLGN spatial integration could be extended by considering the time course of the effects. Indeed, both in the modeling and in the experimental part of our study, we focused on time-averaged responses. We made this choice because L6CT feedback is known to have a wide range of axonal conduction latencies, including very short ones^124^, with feedback effects arriving in dLGN while its feedforward response is still unfolding. Since latencies might not only vary between different types of L6CT neurons^124^, but could also be subject to global trial-by-trial, state-dependent modulations^125^, the most powerful approach to tackling this question would be simultaneous dual- or multi-area recordings within the thalamo-cortico-thalamic loop.

### Manipulating corticothalamic feedback

To probe the effects of CT feedback, we suppressed cortical activity by optogenetic activation of V1 PV+ inhibitory interneurons. Because it relies on intracortical inhibition, this method provides strong suppression throughout all layers of cortex^53^. However, its global character limits the specificity with which individual circuits can be targeted^126^. In the case of CT feedback, global V1 suppression might not only modulate thalamic activity via the L6CT circuit, but also via other, polysynaptic pathways. One potential alternative route could arise from corticofugal neurons in layer 5 that are known to influence the gain of responses in the superior colliculus (SC)^127, 128^, which in turn provides excitatory, driving input to dLGN^129^. We regard it as unlikely that any of the effects observed in our study are mediated via the SC. First, effects of direct SC suppression on dLGN responses are limited to the dorsal-most 150 *µ*m of the dLGN^130^, while effects evoked by V1 suppression in our study spanned a substantial depth. Secondly, suppressing V1 affects SC responses independently of stimulus size^128^, which is inconsistent with the size-dependent effects we found for dLGN. Other alternative influences could involve undesired rebound effects at the edge of the cortical area undergoing optogenetic suppression. We think it is unlikely that intracortical rebound effects drive our main conclusions, because, firstly, such effects have not been prominent in previous studies quantifying the lateral extent of cortical suppression^53, 131^, and secondly, we have found a consistent spatial pattern of CT feedback effects in dLGN with both global V1 suppression and L6CT photoactivation, which likely recruit intracortical circuits in different ways. To rule out the effects of polysynaptic circuits during global suppression, it is not sufficient to selectively suppress L6CT pyramidal cells at the level of V1^34, 132^, because they have an intracortical axon collateral that targets layer 5^50^, and also make privileged connections with a translaminar PV+ interneuron subtype in L6^51, 52^, which in turn strongly regulates the gain of the entire V1 column^34, 51, 52^. Instead, a more promising next step would be to directly suppress axon terminals of L6CT pyramidal cells at the thalamic target. This is challenging, because direct optogenetic inhibition of axon terminals is prone to unintended excitatory effects^126, 133^, due to changes in pH in the case of light-driven proton pumps^134^, and a depolarizing reversal potential for chloride in the case of anion-selective channelrhodopsins (ACRs)^126^.

Our results contribute to an emerging view, which posits that manipulation of L6CT pyramidal cells does not simply produce global gain changes in dLGN, and that photostimulation and photosuppression do not simply produce changes with opposite sign. First, effects of L6CT activation, as shown here and consistent with previous studies^20, 135^, cannot be described by a global gain factor, because these effects have a spatial profile, ranging from a combination of suppression and facilitation in the dLGN region corresponding to the retinotopic location of the L6 source to suppression beyond. Second, effects of CT feedback on sensory thalamic nuclei are known to be frequency-dependent^19, 136^, reflecting the distinct short-term dynamics of synapses in the CT circuit. In particular, high-frequency CT feedback stimulation has been reported to yield net excitatory effects, due to the facilitation of the L6CT pyramidal cell synapses and depression of the TRN synapse, while low-frequency CT feedback stimulation seems net suppressive^19^. Complicating the matter further, CT feedback can increase dLGN firing not only via net depolarization, but also by sustained hyperpolarization and rebound firing^64^, through de-inactivation of low-threshold, T-type Ca^2+^ channels^56^ and subsequent bursting^137^.

L6CT pyramidal cells as targeted by the Ntsr1-Cre line^38, 50–52^ are not homogeneous, but are known to contain at least two subtypes defined by morphology^51, 52, 123, 138, 139^, three subtypes defined by electrophysiology and morphology^139^, and four major subtypes defined by transcriptomics^138, 139^. For the mouse, it is currently unknown whether these subtypes differentially contribute to modulation by CT feedback. In the visual system of primates and carnivores, where feedforward processing along the retino-geniculo-cortical pathway occurs in functionally segregated, parallel pathways, CT feedback circuits seem to mimic this organization, both in terms of morphology^140, 141^ and function^142^ of L6CT pyramidal cells. Functional cell typing of our dLGN and visTRN population only revealed subtle, if any, differential effects of CT feedback, which might not be so surprising given our strategy of global V1 suppression. In the future, investigating to which extent excitatory and inhibitory feedback pathways are recruited under different stimulus and behavioral conditions, and with specificity for L6CT subtypes, will likely yield more complete answers.

## Acknowledgements

This research was supported by DFG BU 1808/5-1 (LB), DFG SFB870 TP19 (LB), by an add-on fellowship from the Joachim Herz Stiftung (GB), by DFG SFB 870 Z04 (M. Götz), and by the Viral Vector Facility of the LMU. We thank A. Wal for recording some of the data in **Figure 2**, and I. Mühlhahn for CsCl DNA preparation. Confocal microscopy was performed in the core facility bioimaging of the LMU Biomedical Center. We are grateful to M. Sotgia for lab management and support with animal handling, M. Sotgia and H. Wolfrohm for contributing to histology, S. Schörnich for IT support, and B. Grothe for providing an excellent research infrastructure.

## Author contributions statement

Conceptualization, L.B., S.E., G.B., F.A.S., A.K.; Methodology, M.H.M., G.E.; Software, G.B., S.E., F.A.S., M.A.S., M.H.M., L.B.; Formal Analysis, G.B., F.A.S., S.E., L.B.; Investigation, G.B., F.A.S., S.E., A.K., M.A.S.; Resources, C.L.L.; Data Curation, M.A.S., G.B., S.E., L.B., F.A.S.; Writing – Original Draft, G.B., S.E., F.A.S., L.B.; Writing – Review & Editing, L.B., S.E., all authors; Visualization, G.B., F.A.S., S.E., L.B.; Supervision, L.B.; Project Administration, L.B.; Funding Acquisition, L.B., G.B.

## Additional information

### Methods

All procedures complied with the European Communities Council Directive 2010/63/EC and the German Law on the Protection of Animals, and were approved by local authorities, following appropriate ethics review. Experiments were performed on three strains of adult mice of both sexes: C57BL/6J (*n* = 3, mean age = 14.2 weeks) and B6;129P2-Pvalb^tm1(cre)Arbr^/J (*n* = 19, mean age = 23.3 weeks), both obtained from the Jackson Laboratory and B6.FVB(Cg)- Tg(Ntsr1-cre)GN220Gsat/Mmcd (*n* = 20, mean age = 24.4 weeks, MMRRC).

#### Virus used for triple-color tracing

pAAV-CAG-FLEX-GFP was a gift from Malin Parmar, pAAV-CAG-FLEX-mScarlet was a gift from Rylan Larsen (Addgene # 99280; http://n2t.net/addgene:99280; RRID:Addgene_99280), and pAAV-TRE-DIO-mTurquoise2 was a gift from Viviana Gradinaru (Addgene # 99115; http://n2t.net/addgene:99115; RRID:Addgene_99115)^S1^. pAAV-CAG-FLEX-mTurquoise2 was generated in the viral vector facility of the LMU by the restriction digest and ligation method, using pAAV-CAG-FLEX-mScarlet as a pAAV backbone, replacing mScarlet with AscI-FseI/Blunt restriction sites, and inserting “mTurquoise2” (AscI/NheI-Blunt) from pAAV-TRE-DIO-mTurquoise2.

#### AAV production

High-titer preparations of rAAV2/1 and rAAV2/retro were produced based on the protocol by Zolotukhin and colleagues^S2^ with minor modifications. In brief, HEK 293T cells (ATCC CRL-3216) were transfected with the CaPO4 precipitate method. For triple-color viral tracing, the pAAV plasmid, Ad helper (Cell Biolabs, Cat.N: gb AF369965.1) and pRC1 (Cell Biolabs) were applied in an equimolar ratio. For retrograde viral tracing, the plasmids rAAV2-retro (Addgene #81070)^72^, Ad helper (Cell Biolabs, Cat.N: gb AF369965.1) and pAAV-CMV-GFP (Cell Biolabs, Cat.N: AAV-400) were applied in an equimolar ratio. All plasmids were purified on CsCl gradients. After 72–96 h, the cell pellet was harvested with the AAV release solution, 50 U/ml benzonase was added, and the solution was then incubated for 2 h at 37° C (water bath). Cells were frozen and thawed in liquid nitrogen to allow rAAV release. Purification of the rAAV vector was done on iodixanol gradients (consisting of 15, 25, 40 and 56% iodixanol), followed by gradient centrifugation at 50,000 rpm for 2 h 17 min at 22° C in a Ti70 rotor (Beckman, Fullerton, CA, USA). rAAV was collected from the 40% iodixanol layer with a 5 ml syringe. rAAVs were dialyzed (Slide-A-Lyzer 10,000 MWCO 5 ml) in buffer A overnight to remove iodixanol. An anion-exchange chromatography column (HiTrap Q FF sepharose) equipped with Superloop was connected with the ÄKTAprime plus chromatography system to collect the eluted fraction. To measure rAAV concentration, the eluted fraction was spun and washed once in PBS-MK Pluronic-F68 buffer with a Millipore 30K MWCO 6 ml filter unit. rAAVs were stored in a glass vial tube in 4° C. rAAVs titers were measured by SYBR Green qPCR with GFP or SV40 or ITR2 primer^S3^. Usual titer was 3 *×* 10^14^ to 5 *×* 10^16^ GC/ml.

#### Surgical procedures for headpost implantation, virus injection and craniotomy

The majority of experiments were performed under Licence ROB-55.2-2532.Vet_02-17-40: thirty minutes prior to surgery, an analgesic (Metamizole, 200 mg/kg, sc, MSD Animal Health, Brussels, Belgium) was administered. Anesthesia was induced by placing the mice in an induction chamber and exposing them to isoflurane (5% in oxygen, CP-Pharma, Burgdorf, Germany). Animals were then fixated in a stereotaxic frame (Drill & Microinjection Robot, Neurostar, Tuebingen, Germany), and the isoflurane level was adjusted (0.5%–2% in oxygen) to maintain an appropriate level of anesthesia, as evaluated by the absence of a pedal reflex. During the procedure, the eyes were protected with an ointment (Bepanthen, Bayer, Leverkusen, Germany) and the animal’s body temperature was maintained at 37° C by means of a closed-loop temperature-control system (ATC 1000, WPI Germany, Berlin, Germany). An additional analgesic was then delivered (Buprenorphine, 0.1 mg/kg, sc, Bayer, Leverkusen, Germany). After the animal’s head had been shaved, the skin was thoroughly disinfected with iodine solution (Braun, Melsungen, Germany), a local analgesic (Lidocaine hydrochloride, 7 mg/kg, sc, bela-pharm, Vechta, Germany) was injected under the scalp, and a small incision was made along the midline. Part of the skin covering the skull was removed, and tissue residues were cleaned by administration of a drop of H_2_O_2_(3%, AppliChem, Darmstadt, Germany). The animal’s head was then adjusted to a skull-flat configuration using four landmarks (bregma, lambda, and two points 2 mm to the right and to the left of the midline, respectively). In mice targeted for head-bar implantation and electrophysiological measurements, OptiBond FL primer and adhesive (Kerr dental, Rastatt, Germany) were applied to the exposed skull, except in locations reserved for subsequent craniotomy, and a site approximately 1.5 mm anterior and 1 mm to the right of bregma, where a miniature reference screw (00-96 X 1/16 stainless steel, Bilaney) soldered to a custom-made connector pin was implanted.

For Cre-dependent triple-color viral tracing, three small craniotomies were drilled over V1 along an iso-azimuth (n = 2) or iso-elevation (n = 2) line or along one coronal section (n = 1) (iso-azimuth: AP:(-3.80, -3.30, -2.80), ML:(-2.70, -2.40, -2.10); iso-elevation: AP:(-3.80, -3.30, -2.80), ML:(-2.60, -2.30, -2.00); coronal section: AP: -3.64, ML:(-2.66, -2.23, -1.79)). 25 nl-50 nl of pAAV-CAG-FLEX-GFP, pAAV-CAG-FLEX-mScarlet, and pAAV-CAG-FLEX-mTurquoise2 (all titres adjusted to 2 *×* 10^15^*/ml* by dilution with a custom-made virus buffer (sterile PBS, 2.6mM KCl, 1mM MgCl_2_, 0.05% Pluronic F68)) were injected into V1 through the three craniotomies, respectively, at a depth of 900 *µ*m. Injections were carried out using a Hamilton syringe (SYR 10 *µ*L 1701 RN no NDL, Hamilton, Bonaduz, Switzerland) equipped with a glass pipette, controlled by the Injection Robot of the Neurostar Stereotax. The craniotomies were covered with sterile bone wax (AngioTech, Vancouver, Canada) and the skin was sutured.

For Cre-dependent expression of ChR2 in PV-Cre and Ntsr1-Cre mice, 2 *µ*L of an adeno-associated virus [pAAV- EF1a-double floxed-hChR2(H134R)-EYFP-WPRE-HGHpA (Addgene, #20298), with different serotypes and titers *≥* 7 *×* 10^12^ vg/mL)] was mixed with 0.3 *µ*L fast green (Sigma-Aldrich, St. Louis, USA). A small craniotomy was performed over V1 at (AP: *−*2.8 mm, ML: *−*2.5 mm), (AP: *−*2.8 mm, ML: *−*2.3 mm), (AP: *−*3.08 mm, ML: *−*2.5 mm) or (AP: *−*3.28, ML: *−*2.4 mm) to enable injection of the prepared mixture. In PV-Cre mice, a total of *∼* 0.2 *−* 0.5 *µ*L of the mixture was injected at multiple depths between 1000 *µ*m and 100 *µ*m below the pial surface. In Ntsr1-Cre mice used for global L6 photostimulation (**Figure S5**), *<* 0.5 *µ*L was injected at depths between 800 *µ*m and 1000 *µ*m, approximately targeting L6. In three Ntsr1-Cre mice used for mapping of L6CT feedback (**Figure 1**), only *∼* 0.05 *µ*L was injected at a depth of *∼* 900 *µ*m. For retrograde labelling of visTRN cells, 0.5 *µ*L of the adeno-associated viral vector rAAV2/retro CMV-GFP (titer: 1.61 *×* 10^15^ GC/ml) was mixed with 1.5 *µ*L PBS and 0.3 *µ*L fast green. In three mice, a small craniotomy was performed above dLGN (AP: *−*2.3 mm, ML: *−*2.3 mm) and 0.4 *µ*L of the prepared mixture was injected at a depth of *−*2.8 mm. Injections were carried out using a Hamilton syringe (SYR 10 *µ*L 1701 RN no NDL, Hamilton, Bonaduz, Switzerland) equipped with a glass pipette, controlled by the Injection Robot of the Neurostar Stereotax.

For all animals planned for electrophysiological recordings (with or without prior virus injection), a custom-made lightweight stainless steel head bar with a cutout for subsequent craniotomy was attached with dental cement (Ivoclar Vivadent, Ellwangen, Germany) above the posterior part of the skull and on top of the primer/adhesive. At the end of the procedure, the cutout was covered with the silicone elastomer sealant Kwik-Cast (WPI Germany, Berlin, Germany). In some animals, an antibiotic ointment (Imex, Merz Pharmaceuticals, Frankfurt, Germany) was applied to the borders of the wound.

For all animals, the long-term analgesic (Meloxicam, 2 mg/kg, sc, Böhringer Ingelheim, Ingelheim, Germany) was injected immediately following the surgery and continued to be administered at 24 h intervals for 3 consecutive days. For a period of 5 days post-surgery, the animal’s health status was assessed with a score sheet.

A smaller number of mice (*n* = 15) were treated in accordance with Licence CIN 4/12, in which general surgical procedures were identical to the foregoing, with the following exceptions. After induction of anesthesia, mice were additionally injected with atropine (Atropine sulfate, 0.3 mg/kg, sc, Braun, Melsungen, Germany). The headpost consisted of a small S-shaped piece of aluminum, which was cemented to the skull between lambda and bregma and to the right of the midline. Virus was injected with either a Picospritzer (Parker Hannifin, Hollis, USA) or a Nanoject (Drummond Scientific, Broomall, USA). Posterior to the head post, overlying the cerebellum, two miniature screws serving as ground and reference were implanted. A well of dental cement was formed over the target recording and stimulation sites and filled with Kwik-Cast. At the end of the procedure, antibiotics (Baytril, 5 mg/kg, sc, Bayer, Leverkusen, Germany) and a long-term analgesic (Carprofen, 5 mg/kg, sc, Rimadyl, Zoetis, Berlin, Germany) were administered and continued to be given for 3 days post-surgery.

To compare visTRN RFs to dLGN RFs (**Figure 4i**), we included dLGN recordings from 16 mice (8 PV-Cre and 8 Ntsr1-Cre mice). In 6 of the Ntsr1-Cre mice, V1 was injected with a virus irrelevant to the purpose of our investigation [AAV-DJ-Ef1a-DIO SwiChR++-EYFP, *n* = 2; pAAV_hSyn1-SIO-stGtACR2-FusionRed (Addgene #105677), *n* = 4].

Gradual habituation of the animal to the experimental condition was begun after at least 7 days of recovery. The habituation phase consisted of 3 days of handling followed by 4 days during which the experimental procedure was simulated. In mice prepared for photostimulation experiments, neural recordings were initiated no sooner than 3 weeks post injection to allow enough time for virus expression. One day before the first recording session, mice were anesthetized in the same way as for the initial surgery. For V1 and dLGN recordings, a craniotomy (ca. 1.5 mm^2^) was performed above V1 and dLGN (AP: *−*2 or *−*2.5 mm, ML: *−*2 mm). For TRN recordings, two smaller craniotomies (ca. 1 mm^2^) were performed over V1 and TRN, respectively (V1: AP: *−*2.8 mm, ML: *−*2.5 mm; TRN: AP: *−*1.25 mm, ML: *−*2.15 mm; or AP: *−*1.25 mm, ML: *−*2.2 mm; or AP: *−*1 mm, ML: *−*2 mm). At the end of the procedure, the craniotomy was re-sealed with Kwik-Cast. To avoid residual drug effects during the recordings, the long-term analgesic Metacam was injected only once at the end of the surgery, unless the mouse showed any sign of distress. Experiments started on the day after craniotomy, and were carried out daily for as long as the electrophysiological signal remained of high quality.

#### Electrophysiological recordings and optogenetic manipulations

Recording sessions were carried out in a secluded chamber that allowed us to perform experiments in the absence of any ambient light source. Animals were head-fixed and positioned on an air-cushioned Styrofoam ball that enabled the mouse to move freely. Ball movements were recorded at 90 Hz by two optical computer mice connected to a microcontroller (Arduino Duemilanove). Eye position and pupil size were recorded under infrared light illumination with a Guppy AVT camera (frame rate 50 Hz, Allied Vision, Exton, USA) interfaced with a zoom lens (Navitar Zoom 6000, Rochester, USA). Extracellular activity was sampled at 30 kHz (Blackrock microsystems, Salt Lake City, USA). At the beginning of each recording session, the silicone plug covering the craniotomy was removed and a silicon probe (A1x32Edge-5mm-20-177-A32, A1x32-5mm- 25-177, A1x16-3mm-50-177-A16, A1x64-Poly2-6mm-23s-160, NeuroNexus, Ann Arbor, USA; H3, Cambridge NeuroTech, Cambridge, UK) was positioned above the target site with a micromanipulator (MP-225, Sutter Instrument, Novato, CA, USA) and inserted to the appropriate depth (mean recording depth in *µ*m: V1: 1040; dLGN: 3100; visTRN: 3394) until we encountered vigorous responses to visual stimuli. For recordings from dLGN and TRN, we judged the correct position of the electrode based on *post mortem* histological reconstruction of the electrode track, for which the electrode was stained with a lipophilic fluorescent tracer (DiI, DiD, Invitrogen, Carlsbad, USA) on one of the final recording sessions. For recordings from dLGN, where physiological properties are well known^39,S4^, additional indicators were the characteristic progression of RFs from upper to lower visual field along the electrode shank, the neurons’ preference for drifting gratings of high temporal frequency, and the manifestation of this frequency in the response pattern of the cells (strong F1 response).

To photostimulate PV+ inhibitory interneurons or L6CT cells, we interfaced an optic fiber (910 *µ*m diameter, Thorlabs, Newton, USA; or 480 *µ*m, Doric Lenses, Quebec, Canada) with a blue light-emitting diode (LED) (center wavelength 470 nm, M470F1, Thorlabs, Newton, USA; or center wavelength 465 nm, LEDC2_465/635_SMA, Doric Lenses, Quebec, Canada). The tip of the fiber was placed less than 1 mm above the exposed surface of V1 using a manual micromanipulator. The tip of the head-bar holder was surrounded with black metal foil that prevented the light from reaching the animal’s eyes. For each mouse, the first recording session was conducted in V1 to verify that the photostimulation was effective. Only if light exposure reliably triggered suppression of V1 in PV-Cre mice or activation of L6 in Ntsr1-Cre mice, was the animal used for subsequent recordings from dLGN or TRN. To elicit reliable effects during each recording session, we adjusted the light intensity of the LED on a daily basis (median intensity: 1.1 mW/mm^2^ as measured at the tip of the optic fiber).

#### Visual stimulation

Visual stimuli were presented on a gamma-corrected liquid crystal display (LCD) monitor (Samsung Sync-Master 2233RZ; 47*×*29 cm, 1680*×*1050 resolution at 60 Hz, mean luminance 50 cd/m^2^) positioned at a distance of 25 cm from the animal’s right eye (spanning *∼* 108*×*66°, small angle approximation) and controlled by custom-written software (EXPO, https://sites.google.com/a/nyu.edu/expo/home).

##### RF mapping and identification of cortical layers

We mapped RFs with a sparse noise stimulus, which consisted of non-overlapping black and white squares with a side length of 4 or 5 deg that were arranged on a grid spanning between 40 and 60 deg on each side. Stimulus presentation time varied between experiments and ranged from 0.08 to 0.20 s. Whenever possible, subsequent stimuli were presented at RF locations based on multiunit activity extracted from the ongoing recordings by applying a threshold of 4.5 to 6.5 SD to the high-pass filtered signals.

To determine the V1 laminar location of the recording sites, we presented full-screen, contrast-reversing checkerboards at 100% contrast, with a check size of 25 deg and a temporal frequency of 0.5 cyc/s.

##### Tuning experiments

Drifting gratings adapted in their temporal (0.20 – 15.00 cyc/s) and spatial frequencies (0.01 – 0.08 cyc/deg) to the preferences of neurons at the recording site were used to determine selectivity for orientation, contrast and size. Contrast was set to 1 for all gratings except those in contrast-tuning experiments. In all tuning experiments, we assessed spontaneous firing rate by including trials in which only the mean luminance gray screen was presented. Effects of photostimulation were computed using photostimulation windows and corresponding windows in control conditions during stimulus presentation. Across experiments, we used slight variations of stimulus and light durations. In the figure captions, we indicate the parameters of the most common protocol.

To verify the effectiveness of photostimulation, we performed the first recording session for each animal in area V1, using drifting sinusoidal gratings to measure tuning for various stimulus properties, with photostimulation trials interleaved in pseudorandom order. For the analysis of V1 suppression by photoactivating PV+ inhibitory interneurons (**Figure 2a-c**), we pooled data from direction-tuning experiments (n = 11), size-tuning experiments (n = 19), and contrast-tuning experiments (n = 10). For direction-tuning experiments, grating direction was varied in step sizes of 30 deg or 45 deg. Gratings were presented for 0.75 s with photostimulation starting with stimulus onset and lasting for 0.85 s, or for 1.5 s with photostimulation starting with stimulus onset and lasting for 1.6 s, or for 2 s with photostimulation starting 0.85 s after stimulus onset and lasting for 0.25 s. For size-tuning experiments, gratings ranged in diameter between 0 and 67 deg (in 11 or 15 steps). Stimuli were presented for either 1.5 s with photostimulation starting with stimulus onset and lasting for 1.6 s, or 0.75 s with photostimulation starting 0.21 s after stimulus onset and lasting for 0.25 s. Lastly, for contrast-tuning experiments, contrast was varied in 13 steps between 0 and 1. Stimuli were presented for 2 s, and photostimulation started 0.85 s after stimulus onset and lasted for 0.25 s. For the analysis of L6CT activation effects in V1 during photostimulation of Ntsr1+ neurons (Figure S5a-c), we again pooled data from direction- (n = 11), size- (n = 11), and contrast- (n = 6) tuning experiments. For direction-tuning experiments, grating direction was varied in step sizes of 30 deg. Gratings were presented either for 0.75 s with photostimulation starting 0.1 s before stimulus onset and lasting for 0.85 s, or for 0.75 s with photostimulation starting 0.15 s after stimulus onset and lasting for 0.25 s. For size-tuning experiments, grating diameter was varied between 0 and 67 deg in 13 steps. Gratings were presented for 0.75 s with photostimulation starting 0.10 s before stimulus onset and lasting for 0.85 s, or for 0.75 s with photostimulation starting 0.15 s after stimulus onset and lasting for 0.25 s. Finally, for contrast-tuning experiments, contrast levels were varied between 0 and 1 in 13 steps. Gratings were presented for 0.75 s with photostimulation starting 0.1 s before stimulus onset and lasting for 0.85 s.

To assess the functional specificity of CT feedback (**Figure 1d-i**), we relied on activity measured during orientation-tuning experiments. Sinusoidal gratings drifting in different directions (0 - 330 deg, step size = 30 deg) were presented with and without photostimulation in pseudorandom order. During most experiments (n = 14), stimuli were presented for 0.75 s, and photostimulation started 0.085 s before stimulus onset and lasted for 0.85 s. In a small fraction of experiments (n = 5), stimuli were presented for 1 s, and photostimulation started 0.15 s before stimulus onset and lasted for 1.35 s.

To assess effects of V1 suppression on spatial integration in dLGN (n = 20 experiments, **Figure 2h-m**), we used drifting gratings with stimulus diameter ranging between 0 and 67 deg (in 11 or 15 steps). Gratings were presented for 0.75 s with photostimulation starting 0.04 s before stimulus onset and lasting for 0.85 s, or for 1.5 s and photoactivation starting 0.03 s before stimulus onset and lasting for 1.6 s, or for 0.75 s with photostimulation starting 0.25 s after stimulus onset and lasting for 0.25 s. To probe size tuning in visTRN (n = 69 experiments, **Figure 4j-m**), we used sinusoidal or square-wave drifting gratings with diameters ranging between 0 and 67 deg (in 11 or 15 steps). Stimuli were presented for 0.75 s. In a subset of experiments with paired photoactivation of PV+ neurons in V1 (n = 31, **Figure 5**), photoactivation started 0.04 s before stimulus onset and lasted for 0.85 s.

To measure contrast tuning in visTRN (n = 9 experiments) we presented sinusoidal drifting gratings at different contrasts (in 13 steps). Gratings were presented for 1 s. To measure contrast tuning in dLGN (n = 9 experiments) we presented sinusoidal drifting gratings at different contrasts (in 13 steps). Gratings were presented for 0.75 s (6 experiments) or 0.5 s (3 experiments).

##### Spontaneous activity

To probe the effect of suppressing CT feedback on spontaneous activity in dLGN, we photoactivated PV+ neurons in V1 in the absence of visual stimulation (n = 28 experiments). Photostimulation periods differed between experiments, and ranged from 0.17 s to 1 s.

#### Histology

To verify recording site and virus expression, we performed histological analyses. For experiments under Licence ROB-55.2- 2532.Vet_02-17-40, mice received an analgesic (Metamizole) after the final recording session, and were anesthetized with isoflurane and injected (ip) with a mixture of Medetomidin (Domitor, 0.5 mg/kg, Vetoquinol, Ismaning, Germany), Midazolam (Climasol, 5 mg/kg, Ratiopharm, Ulm, Germany) and Fentanyl (Fentadon, 0.05 mg/kg, Dechra Veterinary Products Deutschland, Aulendorf, Germany) 30 min later. Under deep anesthesia, mice were then perfused with 4% paraformaldehyde (PFA) in phosphate-buffered saline (PBS). Brains were removed, post-fixed in PFA for 24 h, and then rinsed with and stored in PBS at 4° C. Coronal brain slices (40 *µ*m) were cut using a vibratome (Leica VT1200 S, Leica, Wetzlar, Germany), stained with DAPI solution before (DAPI, Thermo Fisher Scientific, Waltham, Massachusetts, USA; Vectashield H-1000, Vector Laboratories, Burlingame, USA) or after mounting them on glass slides (Vectashield DAPI), and cover-slipped. For viral tracing experiments, the perfusion, fixation and slice preparation procedures were identical to those described above, except that brain slices were stained on slide with Invitrogen NeuroTrace Deep-red over night before being cover-slipped.

A scanning fluorescent microscope (BX61 Systems Microscope, Olympus, Tokyo, Japan) was used to inspect slices for the presence of yellow fluorescent protein (eYFP), green fluorescent protein (GFP), and mScarlet (Red fluorescent protein variant), DiI and DiD. Confocal microscopy was performed at the core facility bioimaging of the LMU Biomedical Center with a Leica SP8X WLL microscope, equipped with 405 nm laser, WLL2 laser (470–670 nm) and acousto-optical beam splitter. Images were acquired with a 63x1.30 glycerol objective. For the different fluorophores the following fluorescence settings were used: mTurquoise2 (405; 450–480), GFP (490; 492–550), mScarlet (570; 560–600) and NeuroTrace DeepRed (640; 650–700). Recording was performed in 3 sequences (1: mTurquoise2; 2: GFP, Deep-Red; 3: mScarlet) to avoid bleed through between the channels. All channels were imaged with hybrid photo detectors (HyDs).

For experiments under Licence CIN 4/12, general histological procedures were identical to those described above, except that mice were injected with sodium pentobarbital (Narcoren, 200 mg/kg, ip, Böhringer Ingelheim, Ingelheim, Germany) before perfusion. Coronal brain slices (50 *µ*m) were obtained by using a vibratome (Microm HM 650 V, Thermo Fisher Scientific, Waltham, Massachusetts, USA) and inspected with a Zeiss Imager.Z1m fluorescent microscope (Zeiss, Oberkochen, Germany). For atlas registration and 3D reconstruction, whole brain images were obtained. Images were processed off-line using FIJI^S5, S6^. We adjusted individual color channels for better visibility.

#### 3D reconstruction of expression volumes

For 3D reconstruction and volumetric quantification of expression volumes in L6 and dLGN, brain-slice images had to be annotated and mapped to stereotaxic coordinates for each pixel. To this end, brain-slice images were registered to the Allen Common Coordinate Framework (CCF)^40^, using the allenCCF tools software package (https://github.com/ cortex-lab/allenCCF)^S7^. In brief, for each brain slice, best corresponding atlas sections were chosen manually. To find the optimal transform between atlas coordinates and image pixels, reference points between the atlas section and brain slice image were manually set at unambiguous and salient features of the brain, including structures of the hippocampus, ventricle borders along the midline, habenular nuclei, the midline crossing of the corpus callosum, the indent between the ventral end of the hippocampal formation and the hypothalamus, the meeting point between the medial amygdala and the hypothalamus and high curvature turning points of the brain outline. After successful registration, points set manually along the outline of the expression zones were exported in stereotaxic coordinates. Repeating these steps for the brain slices containing the target regions yielded point clouds in 3D space, circumscribing the expression zones in cortex and thalamus. We computed the convex hull of each point cloud as a geometric description of the expression volume. We chose the convex hull because it is unambiguously defined for any set of points, and does not require prior assumptions about the shape of the volume. To constrain the expression volume with respect to the potentially non-convex structure of the brain area it occupies, we computed the intersection between the convex hull and the 3D model of the brain area of interest (V1 L6 or dLGN). This process yielded a 3D model of that part of the expression zone, which was embedded in the brain area of interest. The intersection operations and computations of volumes on the 3D models were performed with specialized geometry processing software for Python (PyMesh, https://github.com/PyMesh).

#### Locomotion

For recordings under Licence ROB-55.2-2532.Vet_02-17-40 (**Figures 2h-m**, 1i, 4, 5, and associated supplemental figures), we computed run speed by using the Euclidean norm of three perpendicular components of ball velocity (roll, pitch, and yaw)^S8^ and smoothed traces with a Gaussian filter (*σ* = 0.2 *s*). For all analyses of electrophysiological data (except RF mapping with the sparse noise stimulus), we only considered trials in which the animal was sitting. Sitting trials were defined as trials in which the speed of the animal remained below 0.25 cm/s for at least 50% of the time. For recordings performed under Licence CIN 4/12 (Figures 1k–2, and associated supplemental figures), the Gaussian filter differed slightly (*σ* = 0.15 *s*), and hence sitting trials were defined by a run speed below 1 cm/s for 80% of the analyzed time window.

#### Spike sorting

For recordings under protocol ROB-55.2-2532.Vet_02-17-40 (**Figures 2h-m**, 1i, 4, and 5, and associated supplemental figures) were filtered using a 4^th^-order Butterworth high-pass non-causal filter with a low frequency cutoff of 300 Hz. Any saturation in the signal was removed, before clustering responses with the Matlab-based, automated spike-sorting software Kilosort^S9^. The resulting clusters were imported to the Python toolbox Spyke^S10^ for manual refinement of clusters. Spyke allows one to select time ranges and channels around clustered spikes for realignment and for representation in 3D space using dimensionality reduction (multichannel PCA, ICA, and/or spike time). In 3D, clusters were further separated by a gradient-ascent-based clustering algorithm (GAC)^S11^. Using exhaustive pairwise comparison of similar clusters, we merged potentially overclustered units. Only clusters whose autocorrelogram displayed a clear refractory period and whose mean voltage trace showed a characteristic spike waveshape were considered for subsequent analyses.

For data recorded under protocol CIN 4/12 (**Figure 1i**–**Figure 2**, and associated supplemental figures), single neurons in our linear-array recordings were isolated by grouping neighboring channels into 5 equally sized “virtual octrodes” (8 channels per group with 2-channel overlap for 32-channel probes). Using an automatic spike detection threshold^S12^ multiplied by a factor of 1.5, spikes were extracted from the high-pass-filtered continuous signal for each group separately. The first three principal components of each channel were used for semi-automatic isolation of single neurons with KlustaKwik^S13^, and the resulting clusters were manually refined with Klusters^S14^. Only clusters whose autocorrelogram displayed a clear refractory period and whose mean voltage trace showed a characteristic spike waveshape were further considered. In order to avoid duplication of neurons extracted from linear-probe recordings, we computed cross-correlograms (CCHs, 1 ms bins) between pairs of neurons from neighboring groups. Pairs for which the CCH’s zero-bin was 3*×* larger than the mean of non-zero-bins were considered to be in conflict, and only one was kept.

Extracted single units were assigned to the electrode contact with the largest waveform.

#### Analysis of multiunit activity

To obtain robust estimates of RFs at the V1 injection site, we used the envelope of multiunit activity (MUAe), which reflects the number and amplitude of spikes close to the electrode and resembles thresholded multiunit data and average single-unit activity^15, S^^16^. To calculate the MUAe, the median-subtracted, high-pass filtered signals were full-wave rectified, prior to low-pass filtering (200 Hz) and down-sampling to 2000 Hz^S15–S17^.

#### Assignment of units to V1 layers

We assigned units to V1 layers by current-source-density (CSD) analyses^57^. The local field potential (LFP) was computed by downsampling the wideband signal to 1250 Hz. For V1 recordings, the LFP was triggered to contrast reversals of the checkerboard stimulus. The CSD was computed by taking the second spatial derivative of the LFP^57^ and spatially smoothing with a triangular kernel^S18^. The contact closest to the earliest CSD polarity inversion was assigned to the base of layer 4^S19^. The remaining contacts were assigned to putative layers based on a cortical thickness of 1 mm and anatomical measurements of relative layer thickness in mouse V1^143^. Note that the depth estimation is limited in resolution to the electrode site spacing (either 25 or 20 *µ*m, depending on probe configuration), both in terms of estimating the base of L4 as well as taking the contact with largest waveform.

#### Estimation of anatomical depth in the dLGN

To estimate the anatomical depth of recorded neurons in the dLGN, we considered multiunit activity. The top most channel which showed a clear MUAe RF and was well aligned with the characteristic progression of RFs in the dLGN along the dorso-ventral axis^39^ was set as the reference channel estimating the dorsal edge of dLGN during the respective recording session. Single neurons were then assigned the relative depth of the channel with the maximum amplitude of their extracellular waveshape with respect to the reference channel, as determined by the spatial layout of the probe.

#### Data analysis

All further analyses were conducted with custom-written code in Matlab or Python, using the DataJoint framework^S20^. All statistical tests were two-sided.

We calculated mean percent change as

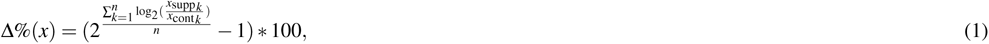

where *x*_supp_ and *x*_cont_ represent the measured variable under the control condition and under the photostimulation condition respectively, and *n* is the number of observations.

##### Descriptive modelling of tuning curves

To characterize neural selectivity, we fit descriptive models and determined goodness-of-fit by *R*^2^ = 1 *−* (*SSE/SST*), where *SSE*=∑(*y−ŷ*)^2^ and *SST*= ∑(*y− ȳ*)^2^.

##### Receptive-field fitting

Receptive-field maps obtained in sparse-noise experiments were fit with a 2D Gaussian^S21^.

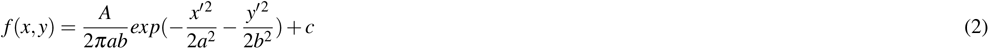

where *A* is the maximum amplitude, *a* and *b* are half-axes of the ellipse, and *x^t^* and *y^t^* are the transformations of the stimulus coordinates *x* and *y*, considering the angle *θ* and the coordinates of the center (*xc, yc*) of the ellipse, and *c* is an offset. RF area (**Figure 4d,h,i** and **Figure 5**) was calculated at 1 *σ* .

In analyses where we relied on MUAe activity (**Figure 1f-i** and **Figure 2h-m**), the RF maps were based on MUAe activity between 50 and 100 ms after stimulus onset (both black and white squares). For the comparison of classical RF sizes in dLGN and visTRN (**Figure 4d,h,i**), the RF maps were based on single-unit responses to both bright and dark stimuli. Before fitting the 2D Gaussian, mean responses were normalized by first subtracting the minimum response and then dividing by the range. Responses in direction-tuning experiments (**Figure 1i** and **Figure 2b**) were fit with a sum of two Gaussians with peaks 180 deg apart, which could have different amplitudes but equal width and a constant baseline^S22^:

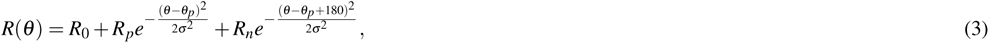

where *θ* is stimulus direction (0–360 deg). The function has five parameters: preferred direction *θ_p_*, tuning width *σ*, baseline response *R*_0_, response at the preferred direction *R_p_*, and response at the null direction *R_n_*.

##### Orientation and direction selectivity

Orientation selectivity was quantified according to^34^,^S23^ as

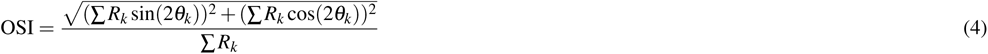

where *R_k_* is the response to the *k*th direction given by *θ_k_*. We determined OSI for each unit during control conditions without optogenetic manipulation.

Direction selectivity index (DSI,^144^) was computed for each unit as

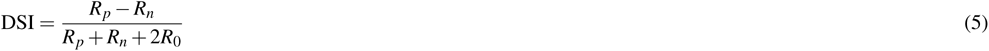

where *R_p_* and *R_n_* are the firing rates in the preferred and null directions, respectively, taken from tuning curves fit to responses to drifting gratings in different directions, and *R*_0_ is baseline firing rate independent of orientation. For both OSI and DSI we focused on experiments in which responses were sufficiently well fit (*R*^2^ *>* 0.8).

##### Contrast sensitivity

We fit contrast response functions with a hyperbolic ratio function^S24^:

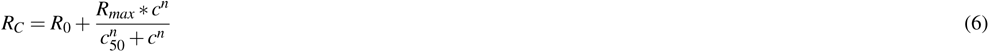

where *c* is the stimulus contrast. The function has four parameters: baseline response *R*_0_, responsiveness *R_max_*, semisaturation contrast *c*_50_, and exponent *n*. To compute contrast response functions we only considered trials in which the animals were sitting. For the analyses we focused on experiments in which the response pattern was well captured by the model (*R*^2^ *>* 0.8).

##### Size tuning

To analyze size tuning in dLGN, we fit responses to drifting gratings of different sizes with a ratio-of-Gaussians model^58^, where a center Gaussian is normalized by a Gaussian representing the surround, each having their independent amplitude (*k*) and width (*w*):

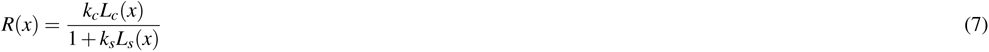

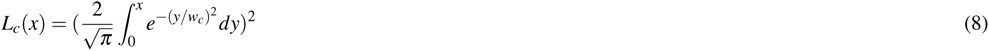

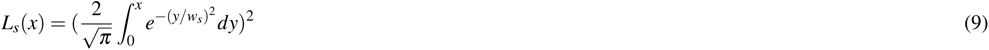

We always constrained *w_c_ < w_s_*.

To analyze spatial integration in visTRN (**Figure 4j-m** and **Figure 5**), we included an offset (*b*) and allowed for rectification of the size-tuning curve to better capture spatial integration in neurons whose firing rates were substantially reduced during V1 suppression:

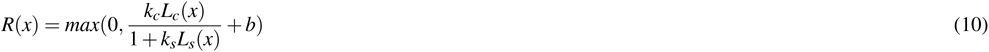

We subtracted the modelled response to stimulus size 0 deg from the resulting curve and quantified suppression strength with a suppression index: *SI* = (*R*_opt_ *− R*_supp_)*/R*_opt_, where *R*_opt_ is the peak response and *R*_supp_ is the response to the largest

stimulus diameter (75 deg). The peak response was defined as the response to the stimulus diameter for which a 1-deg increment in size failed to increase the modelled firing rate by 0.05%. Similar to previous observations^145–147^, for size tuning curves fitted

to both dLGN and visTRN responses, we found a negative correlation between preferred size and suppression strength (SI; visTRN: *R*^2^ = 0.19, slope: -0.003, *p* = 3.4 *×* 10*^−^*^7^; dLGN: *R*^2^ = 0.19, slope: -0.01, *p* = 0.01, Figure S10d,f).

##### Quantification of RFs for functional mapping of CT feedback

To quantify average RF location at the V1 injection site (**Figure 1f**), we computed an RF map based on MUAe activity for each channel. Channels with poor fits to the 2D Gaussians (*R*^2^ *<* 0.4) were not considered for further analyses. Average V1 RF location was obtained by averaging the center positions over all 2D Gaussians. To quantify the retinotopic distance of dLGN neurons with respect to the V1 injection site, we computed the Euclidean distance between their channels’ MUAe RF center and the retinotopic location of the V1 injection site.

##### Spatial profile of CT feedback

To quantify the spatial profile of CT feedback (**Figure 1i**), we used direction-tuning experiments. We focused on visually driven units, defined by evoked firing rates that differed from spontaneous activity by at least 3.29*×* the standard error of the mean (s.e.m.) for at least one direction, with average firing rates *≥* 0.15 sp/s. We computed for each unit and direction the log2 ratio of firing rates with photoactivation to those under the corresponding control condition, before averaging across directions.

To assess the spatial profile of CT feedback effects in dLGN, we grouped neurons according to their retinotopic distance from the V1 injection zone into overlapping bins (15 deg width, 3.3 deg spacing; average number of units per bin: 66; minimum number of units per bin: 32, except for last bin: 7 units), for which we computed the mean. We estimated the 95% CI of the mean effect per bin by resampling with replacement (1000 iterations). To test for spatial regions with a significant CT feedback effect, we used a cluster-based permutation test^S25^. We grouped all neighbouring bins with the same sign and mean log2 ratios that were significantly different from 0 (0 not within 95% CI) into clusters, and computed the sum of absolute mean log2 ratios within those clusters. We then considered the maximum absolute cluster sum value as the test statistic. These steps were then repeated over 10000 iterations with randomly permuted distance values across all neurons. The p-value was the proportion of random permutations which yielded a cluster sum larger than the one from our original dataset.

Next, we classified single neurons into significantly enhanced, suppressed or unmodulated groups, depending on whether their average log2 ratio was above, below or within the 95% interval of the sampling distribution obtained from permuting the photoactivation labels of trials within directions and recomputing the average log2 ratio across directions (10000 iterations). To test whether the proportions of enhanced, suppressed, or non-modulated neurons depended on retinotopic distance, we counted the numbers of each modulation type within 5-deg bins along the retinotopic distance axis, obtaining a 3 *×* 11 contingency table. Statistical test for non-uniformity was done using an omnibus chi-square test, which was followed by post-hoc chi-square tests for each modulation type.

To test whether significantly enhanced neurons were predominantly present close to the injection site, we again applied a cluster-based permutation approach^S25^. We first calculated the adjusted standardized residuals (ASR), defined as the difference between the observed counts in the contingency table and the expected counts under the null hypothesis, adjusted for the row and column totals. For the enhanced neurons, we grouped neighbouring bins with |ASR*| ≥* 1 for the enhanced neurons into clusters, and computed the sum of |ASR| in those clusters. We then considered the maximum cluster sum value as the test statistic. These steps were then repeated over 100000 iterations with randomly permuted distance values across all neurons. The p-value was the proportion of random permutations which yielded a cluster sum larger than the one from our original dataset.

##### Effects of photostimulation on V1 responses

For the quantification of effects of optogenetic manipulations on V1 responses, we only considered V1 neurons whose maximal firing rate exceeded 0.5 sp/s in tuning experiments involving either different directions, sizes or contrasts. Furthermore, we excluded neurons that showed a change in the sign of the effect of optogenetic manipulation across experiments. We first computed, for each unit and experiment, average firing rates during photostimulation in trials with optogenetic manipulation, and in equivalent time windows in trials of the control condition. We then computed, across experiments, the effect of photostimulation by taking the difference in average rates between the photostimulation condition and the control condition, normalized to the rate in the control condition. For the analysis of average effects of V1 suppression by optogenetic activation of PV+ inhibitory interneurons, we excluded putative PV+ inhibitory interneurons directly driven by the light, defined as *≥* 2-fold increase of firing rates in the photostimulation condition compared to the control condition.

##### Effects of V1 suppression on dLGN responses

To analyze effects of V1 suppression on dLGN responses, we considered neurons to be located in dLGN (as opposed to e.g. in the dorsally located hippocampus), if their highest-amplitude extracellular spike waveshape was measured on an electrode channel including and between channels delineating the top and bottom of dLGN. Top and bottom dLGN channels were defined as the dorsal- and ventralmost channels, respectively, with visually responsive neurons in at least one tuning experiment, involving gratings of either different directions, sizes, temporal frequencies or contrasts. We defined a neuron as being visually responsive in these tuning experiments if (1) the absolute difference between its mean firing rates under at least 3 conditions within an experiment and the interleaved blank condition was larger than 2.58*×* the standard error of the mean rate under that condition, and (2) its maximal firing rate exceeded 0.5 sp/s.

For the analysis of effects of V1 photostimulation on dLGN responses to medium gray screen (corresponding to a size 0 deg stimulus, **Figure 2d-g**), we excluded neurons that never spiked in a time window around V1 photostimulation (*±*(0.8 s+Δ*t_opto_*)), where Δ*t_opto_* is the duration of V1 photostimulation. We focused on experiments with a minimum of 5 trials, during which the animal was sitting during the temporal analysis windows of interest. To assess changes in firing rate, we computed for each unit an average firing rate during the window of V1 photostimulation and during a window of equivalent length immediately preceding light onset. For the analysis of burst ratios, we excluded all neurons that did not spike either in the control or the photostimulation window, as the ratio of burst spikes to all spikes in such cases is not defined. We assessed changes in bursting by computing in the same time windows the ratio of burst spikes to the total number of spikes. Burst spikes were defined according to^135^, and required a silent period of at least 100 ms before the first spike in a burst, followed by a second spike with an interspike interval *<* 4 ms. Any subsequent spikes with preceding interspike intervals *<* 4 ms were also considered to be part of the burst. All other spikes were regarded as tonic.

For the analysis of V1 suppression effects on dLGN spatial integration (**Figure 2h-m**), we considered neurons for further analysis whose size-tuning curves had an *R*^2^ *≥* 0.7 and, under the control condition, a mean firing rate of at least 0.15 sp/s. We discarded experiments in which the stimulus center was placed outside of 1 *σ* of its fitted RF center. We focused on RF fits with *R*^2^ *≥* 0.4 obtained from units that responded to the sparse noise stimulus with a sufficiently high firing rate (*≥* 0.15 sp/s). If none of the fitted single unit RFs fulfilled these criteria, we used, if they were well fit (*R*^2^ *≥* 0.4), RFs computed from MUAe activity. To further assure that neurons were well driven by our stimulus, we only included neurons whose mean response to the first 4 stimulus sizes greater than 0 deg was, under the control condition, at least 5% larger than the response to the blank screen. To evaluate the effects of CT feedback on small stimulus sizes, we considered, for each neuron, the responses obtained from the model for the preferred size under the control condition. To assess the effect of CT feedback for large stimulus sizes, we compared modelled responses to a 200 deg stimulus under the two conditions.

##### RFs in visTRN and dLGN

To analyze the organization of classical RFs in visTRN (**Figure 4e-g**) and to compare their sizes to those of dLGN neurons (**Figure 4d,h,i**), we analyzed responses to sparse noise stimuli. We focused on units with a mean firing rate of at least 0.15 sp/s, and whose RFs were well-fit (*R*^2^ *≥* 0.65 for retinotopy in visTRN and *R*^2^ *≥* 0.4 for comparing RF sizes). If for a given unit, results from more than one sparse-noise experiment fulfilled these criteria, we selected the experiment in which the RF was best captured by the 2D Gaussian (largest *R*^2^ value). To test whether the relation between estimated depth and RF center position was different for azimuth versus elevation, we performed an analysis of covariance (ANCOVA), which regressed depth on visual angle and included the categorical covariate azimuth vs. elevation.

##### Spatial integration in visTRN

To analyze spatial integration in visTRN (**Figure 4j-m** and **Figure 5**), we only considered units whose mean firing rate in the control condition was sufficiently high (*≥* 0.15 sp/s) and whose size-tuning curve in the control condition was well captured by the model (*R*^2^ *≥* 0.7). We further concentrated on experiments in which the stimulus center had been presented inside 1 *σ* of the fitted RF center, focusing on RF fits with *R*^2^ *≥* 0.4 obtained from units with sufficiently high mean firing rates (*≥* 0.15 sp/s). In cases where a unit fulfilled these criteria for multiple size-tuning experiments, we focused on the experiment in which responses in the control condition were best captured by the ratio of Gaussians model (largest *R*^2^ value).

Suppression index and preferred size were computed as described above. For few units, our definition of the preferred size and the absence of surround suppression led to slightly stronger responses to the largest stimulus as compared to the optimal stimulus diameter, resulting in negative suppression indices. In such cases, we set the suppression index to 0.

To rule out that a lack of surround suppression could be explained by the difference between stimulus center and RF center or the difference between monitor center and RF center, we computed linear regressions between the suppression indices and the two differences (**Figure S10**). When multiple valid RF mapping experiments were available for a unit, we used the RF with the best model fit (largest *R*^2^ value).

##### Quantifying effects of V1 suppression on visTRN responses

For the analysis of burst ratios (**Figure 5f**), we computed, separately for the control condition and the V1 suppression condition, the ratios of burst spikes to the total number of spikes during sitting trials of a size-tuning experiment. Burst spikes were defined according to^S26^, and required a silent period of at least 70 ms before the first spike in a burst, followed by a second spike with an interspike interval *<* 10 ms. Any subsequent spikes with preceding interspike intervals *<* 10 ms were also considered to be part of the burst. All other spikes were regarded as tonic.

To ensure that suppression indices and preferred size for size-tuning curves recorded under V1 suppression could be reliably interpreted, we required a minimum mean firing rate of 0.1 sp/s during V1 suppression for the analyses in **Figure 5g-h**. Prior to computing population size-tuning curves (**Figure 5d**), differences in response rate as a function of stimulus size (**Figure 5i**), and fitting the threshold-linear model (**Figure 5j-k**), we normalized the fitted size-tuning curves by dividing them by the maximum response across the two conditions.

To analyze differences in response rate between control and photostimulation conditions as a function of stimulus size (**Figure 5i**), we subtracted for each unit the normalized size-tuning curve (1 deg resolution) in the control condition from that in the photostimulation condition, and took the mean across the population. To test for a significant change in the effect of photostimulation with size, we computed the difference in photostimulation effect for subsequent sizes (1 deg steps) and used a resampling procedure across neurons (1000 iterations). If 0 was outside the 97.5^th^ percentile of the resulting distribution of mean differences, we considered the change significant.

To characterize the change in visTRN size tuning induced by suppression of CT feedback (**Figure 5j-k**), we predicted visTRN responses to stimuli of different sizes during V1 suppression based on responses under the control condition by fitting a threshold-linear model:

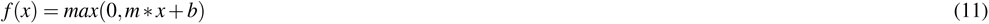

If the resulting fit was of good quality (*R*^2^ *≥* 0.8), we extracted the slope and the threshold parameter (x-intercept).

#### Computational modeling

To explore how dLGN size-tuning changes with the spatial scale of the inhibitory CT feedback component, we used the extended difference-of-Gaussians (eDOG) model^59^. This choice of model was motivated by our intention to explore the effects of CT feedback in a conceptually and mathematically simple framework. The eDOG model is a mechanistic firing-rate based model, in which visual responses of dLGN relay cells are computed from direct evaluation of integrals, representing the spatiotemporal receptive field of RGCs, and feedforward and feedback coupling kernels connecting the neurons of the circuit. Despite the relatively simple linear mathematical structure of the model, it nevertheless incorporates two key biological features of the CT feedback: (i) the non-linearity (half-wave rectification) of cortical L6 cells providing feedback and (ii) the observation that dLGN cells receive feedback from numerous cortical cells with different orientation selectivities together covering all directions. A key simplification of the framework is that the different cortical populations providing CT feedback can be modelled to be uncoupled at the cortical stage. Thus, the eDOG model does not explicitly consider intracortical connectivity or transformations, such as between the dominant input layer 4 and layer 6, from where the CT feedback arises. Rather, such effects are implicitly contained in the choice of effective spatiotemporal feedback coupling kernels.

To carry out simulations in the eDOG modeling framework, we computed response curves using the python toolbox pyLGN^60^ (**Figure 3**). We evaluated the model in its mixed-feedback configuration, where a given dLGN relay cell receives feedback of both signs from cortical cells belonging to the ON and OFF pathways. We took existing code (https://github.com/miladh/edog-simulations/tree/master/size_tuning) that had specified the model parameters following insights from the cat visual system and adjusted them to mimic more closely the properties of the mouse visual system. For the difference-of-Gaussians which represents the receptive field of retinal ganglion cells (RGCs), we approximated width parameters based on data recorded from transient OFF-*α* RGCs^S27^. For the coupling kernels, we scaled the width parameter by a factor of 10, excluding the target inhibitory feedback kernel which we varied between 1 and 40 in 1-deg steps. For each inhibitory feedback kernel width, we then generated tuning curves by simulating responses to static gratings of different size (diameter = 0 - 75; stepsize = 1 deg) with and without feedback. Feedback was manipulated by setting the weight of the feedback kernels to either 0 (no-feedback) or 1. The resulting curves were normalized so that the maximum response in the no-feedback condition equaled 1. Preferred size and suppression index were computed as described for the electrophysiological data.

#### Data and code availability

Except for **Figure 3**, all figures were generated from processed data. The data and code used to generate the figures will be made available upon acceptance of the manuscript.

**Figure S1.**
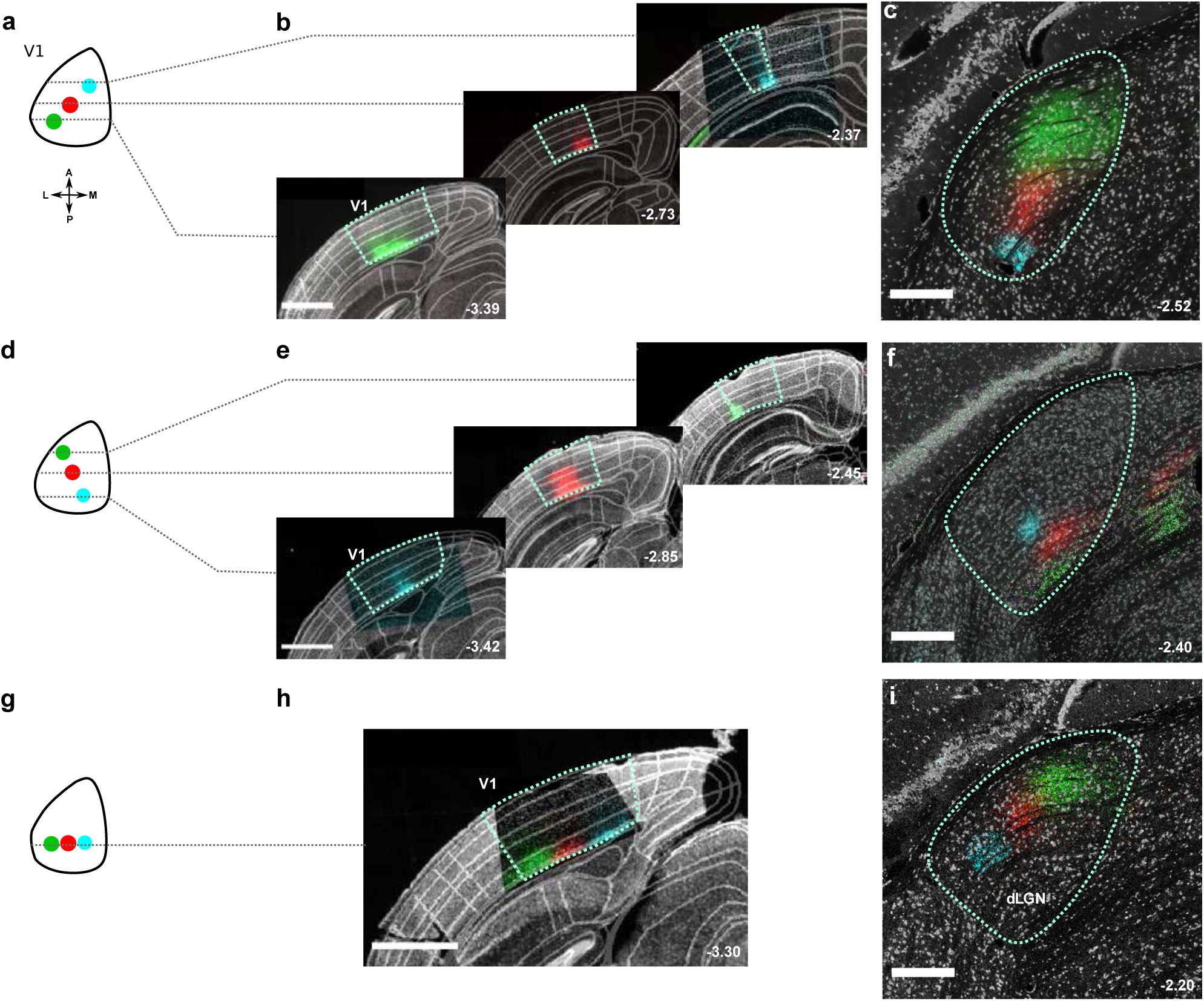
Results of additional mice used for the analysis of retinotopy of corticothalamic projections. (**a**) Cartoon of V1 injection sites along the elevation axis. (**b**) Representative coronal slices with fluorophore expression along the V1 elevation axis. Images are ordered posterior to anterior. (**c**) Labeled L6CT axonal terminal fields in dLGN. (**d–f**). Same as (a–c) for another mouse, injected along the V1 azimuth axis. (**g–i**) Results of another mouse, where V1 injections were placed within a single coronal plane. Narrow-field images of *mTurquoise2* in b,e and *eGFP, mScarlet* and *mTurquoise2* in h were acquired with a confocal microscope and manually aligned with the wide-field epifluorescence images of the corresponding brain slices. All panels: small numbers indicate distance from bregma. (b,e,h) Scale bar: 1 mm. (c,f,h) Scale bar: 250 *µ*m

**Figure S2.**
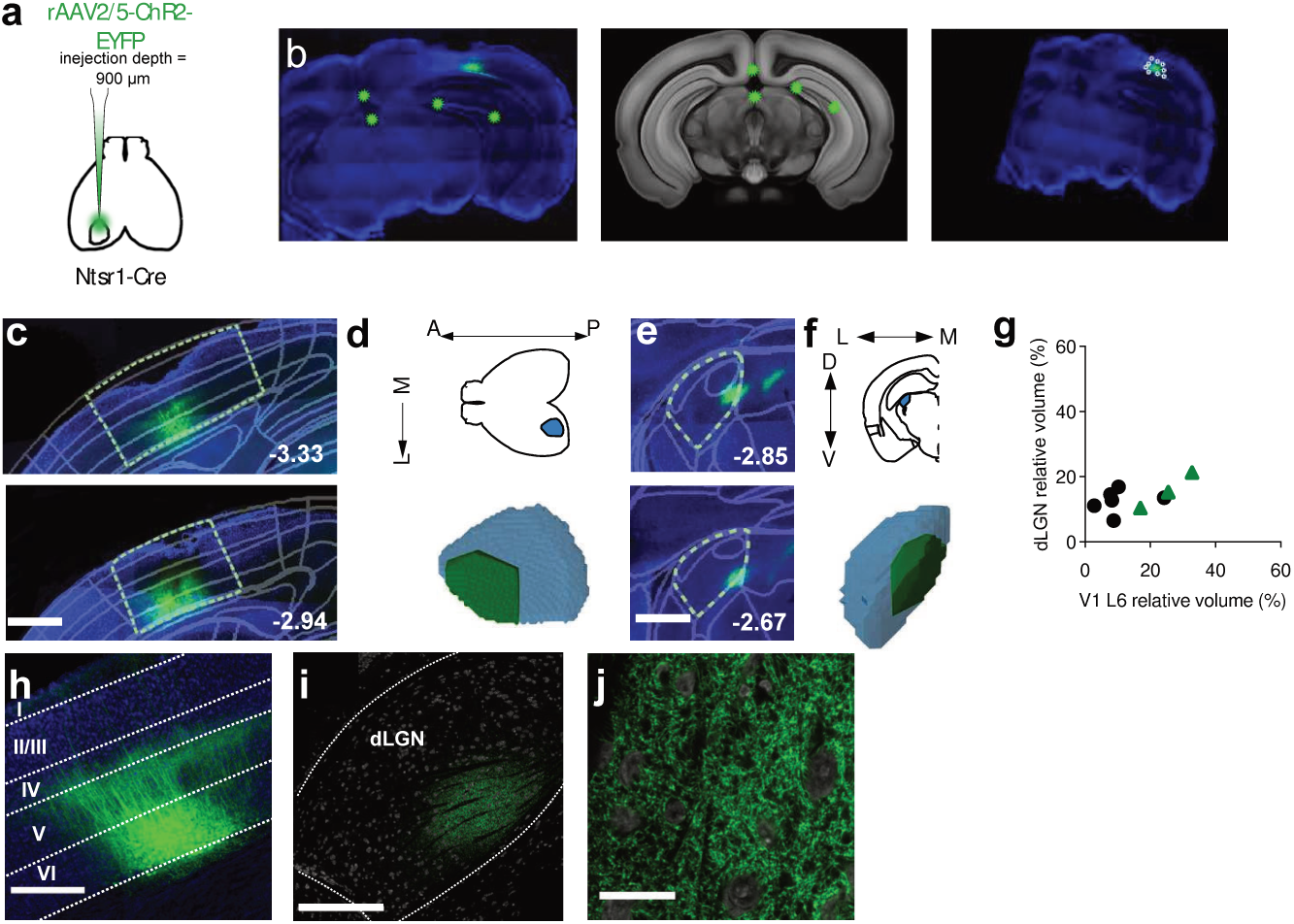
Quantification of expression volumes for ChR2-assisted functional mapping and viral tracing of CT feed- back through registration of the *post-mortem* histological data from individual mice into a standardized 3D anatomical coordinate system (Allen Mouse Common Coordinate Framework [CCF]^40^) and quantification of relative expression volumes. (**a**) Schematic of the experiment: To examine the spatial specificity of CT feedback in mouse dLGN, we expressed ChR2-eYFP (n=3), eGFP (n=3) or mScarlet (n=3) in a localized population of V1 L6CT pyramidal cells, by injecting a small volume of Cre-dependent AAV into V1 in Ntsr1-Cre mice^38^ (see also Figure 1a**,b** and **Figure S1**). (**b**) Pipeline for quantification of expression volumes. (b) *Left*: Manually chosen reference points (*green circles*) on salient features of an example brain-slice image. *Blue*: DAPI; *fainter green*: eYFP. *Middle*: Corresponding locations of the reference points are marked on the manually chosen atlas section from the Allen CCF^40^. (c) *Right*: Brain-slice image registered and transformed to the CCF. White points outline the expression zone and are extracted as CCF coordinates. (**c–f**) Computation of the relative volumes of transduced V1 CT pyramidal cells within L6 (“source volume”) and those of their dLGN projections (“target volume”) for a representative Ntsr1-Cre mouse. (c) Coronal sections of the area close to the V1 injection site are overlaid with fitted area boundaries from the Allen CCF (*gray*). *Green*: ChR2-eYFP. (d) *Top*: Axial schematic of V1 L6 (*blue*) within the cortex (*black contour*). *Bottom*: 3D reconstruction of the expression volume (*green*) within V1 L6 (*blue*), seen from the same perspective as the upper panel (“source volume”). Relative volume: 25%. (e) Coronal sections showing that axons of the transduced L6CT neurons project to a restricted volume the dLGN. (f) *Top*: Coronal schematic of dLGN (*blue*) within the brain section (*black contour*). *Bottom*: 3D reconstruction of the expression volume (*green*) within the dLGN (*blue*), seen from the same perspective as the upper panel (“target volume”). Relative volume: 15%. In (c,e), numbers in bottom right corner indicate distance from bregma in mm; scale bar 0.5 mm. (**g**) Comparison of the relative expression volumes within V1 L6 ((expression volume within V1 L6)/(total volume of V1 L6)) and dLGN ((expression volume within dLGN)/(total volume of dLGN)) for each mouse. Local injections in V1 yield restricted, spatially specific expression in dLGN with similar relative volumes (mean difference = 0.017, p = 0.55, resampling). *Black*: mice used for viral tracing experiments; *green*: mice used for ChR2-assisted functional mapping). Local injections in V1 yield restricted, spatially specific expression in dLGN. (**h**) Example close-up image of L6 CT neurons expressing eGFP (*Green*). Scale bar: 0.5 mm. (**i**) Example confocal image of dLGN with eGFP signal in projections from L6 CT neurons. Scale bar 250 *µ*m. (**j**) Close-up confocal image of L6 CT projections in dLGN in same slice as in i. Scale bar 25 *µ*m.

**Figure S3.**
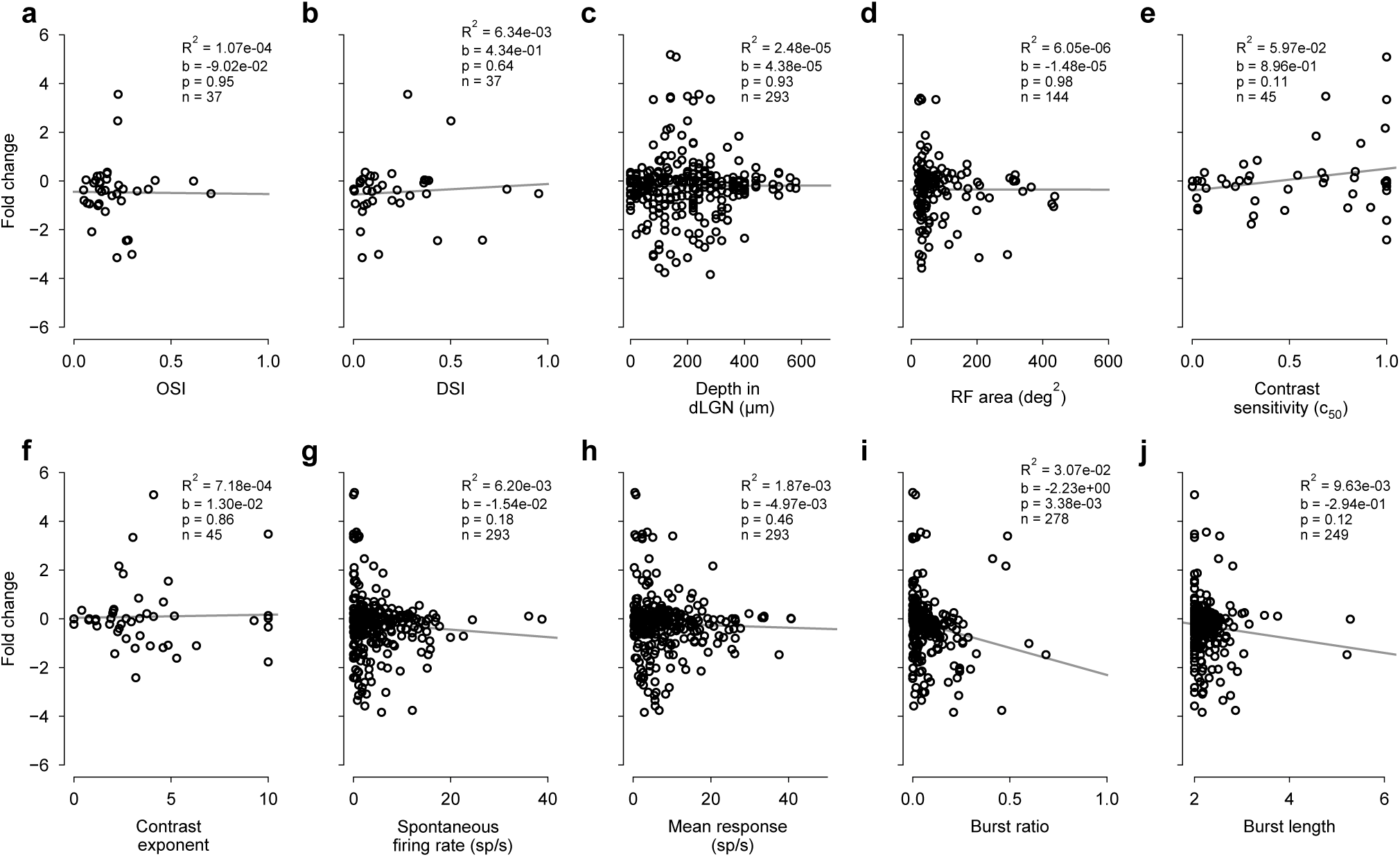
Diversity of effects of V1 L6CT photoactivation on dLGN firing rates cannot be explained by functional response properties. Change of firing rate (fold change, log_2_ ratio) as a function of (**a**) orientation selectivity index (OSI^34^), (**b**) direction selectivity index (DSI)^144^, (**c**) estimated depth within dLGN, (**d**) RF area as obtained from sparse noise experiments, (**e**) contrast sensitivity (*c*_50_) and (**f**) exponent *n* of the contrast response function, (**g**) spontaneous firing rate obtained from interleaved blank trials, (**h**) mean response across all drift directions, (**i**) burst ratio, and (**j**) burst length (spikes/burst). Functional properties in (a,b,g–j) are computed from direction selectivity experiments. *b*: slope; *p*: significance of slope.

**Figure S4.**
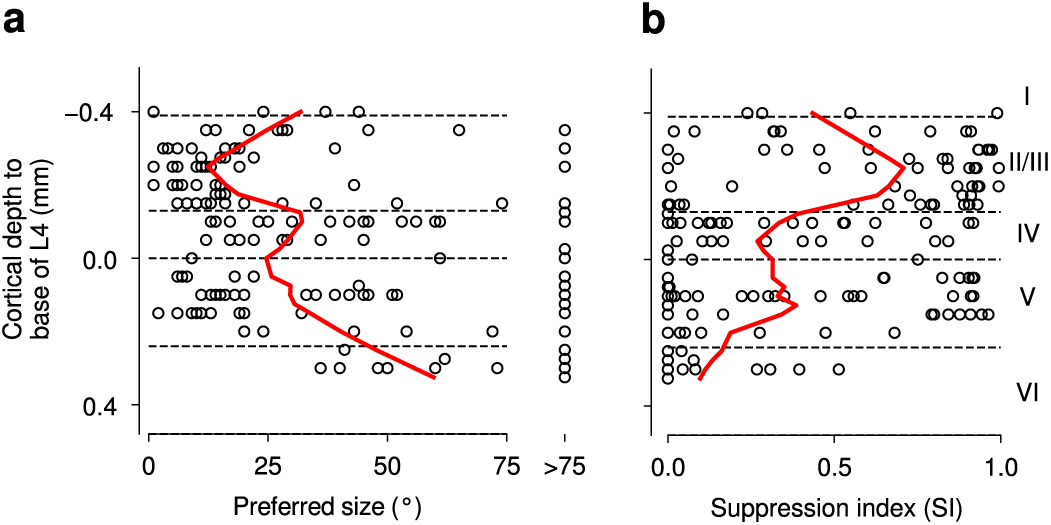
Neurons in L6 of mouse V1 prefer large stimulus sizes and experience little surround suppression. (**a**) Distribution of preferred size for neurons (n = 177) recorded across layers of V1. (**b**) Same as (a) for suppression index. *Dashed horizontal lines*: borders between V1 layers, based on CSD analysis^57^ and histological estimates of relative layer thickness^143^. *Red*: Smoothed mean computed by local robust regression (MATLAB funtion “smooth”, method “rlowess”, window size = 0.28 mm).

**Figure S5.**
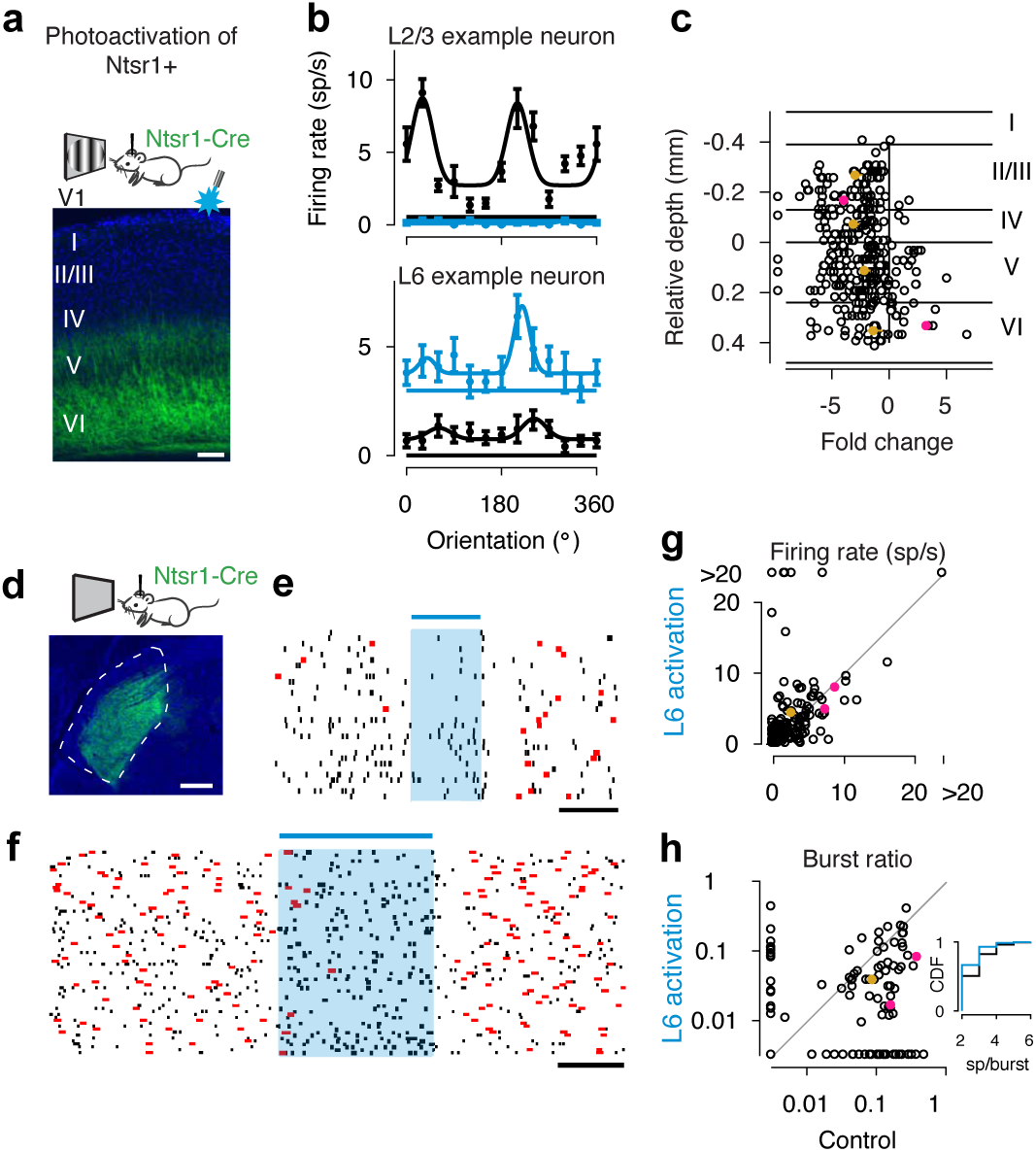
Photoactivation of L6CT neurons promotes dLGN tonic firing mode. (**a**) Coronal section of V1 of a Ntsr1-Cre mouse injected with Cre-dependent AAV-ChR2. *Green*: ChR2-YFP, *blue*: DAPI. Scale bar 100 *µ*m. (**b**) Example orientation-tuning curves of cells located in putative L2/3 or putative L6 for trials during V1 L6CT photoactivation (*blue*) and under control conditions (*black*). Drifting gratings with temporal and spatial frequencies coarsely optimized for the recording were presented for 0.75 s with continuous photostimulation starting 0.1 s before stimulus onset and lasting for 0.85 s. (**c**) Fold change (i.e. log2 ratio of average firing rates for V1 L6CT photoactivation and control conditions across tuning experiments) as a function of cortical depth relative to the base of L4, estimated by CSD^57^. Consistent with previous literature, Ntsr1+ neurons seem to provide a mix of both excitation and di-synaptic inhibition in lower layers, while exerting mostly di-synaptic inhibition in upper layers^34, 50, 51^. *Gold*: layer-wise mean; *pink*: example neurons. Error bars: confidence intervals of the mean, determined by bootstrapping. n = 362 neurons. (**d**) Coronal slice of dLGN, with axons of Ntsr1+ neurons expressing ChR2 in *green*. (**e-f**) Recordings from dLGN. Raster plots of two example dLGN neurons during spontaneous activity aligned to V1 L6CT photoactivation (*shaded blue*). *Red*: burst spikes, black horizontal bar: 200 ms. (**e**) n = 31 trials, (**f**) n = 69 trials. (**g**) Firing rates during vs. before V1 L6CT photoactivation. Activation of L6CT neurons did not significantly affect firing rates (during: 4.2 sp/s vs. before: 2.7 sp/s; n = 167 neurons; *p* = 0.4, Wilcoxon signed-rank test), as effects of Ntsr1+ activation on the firing of individual dLGN neurons were diverse. This diversity is consistent with the interpretation of our functional mapping experiments, in which we also observed a combination of enhancement and suppression for dLGN neurons with “nearby” RFs (Figure 1k). (**h**) Ratio of burst spikes during vs. before V1 L6CT photoactivation. Activating CT feedback decreased the fraction of spikes fired in bursts (before: 9.04%, during: 3.75%; n = 139 neurons; *p* = 1.7 10 7, Wilcoxon signed-rank test). Data points at marginals represent neurons whose burst ratio was 0. *Inset*: cumulative distribution of burst lengths during (*blue*) vs. before (*black*) V1 L6CT photoactivation. Activating CT feedback shifted the distribution of spikes per burst towards lower values (*p* = 7.8 *×* 10*^−^*^5^, two-sample Kolmogorov-Smirnov test).

**Figure S6.**
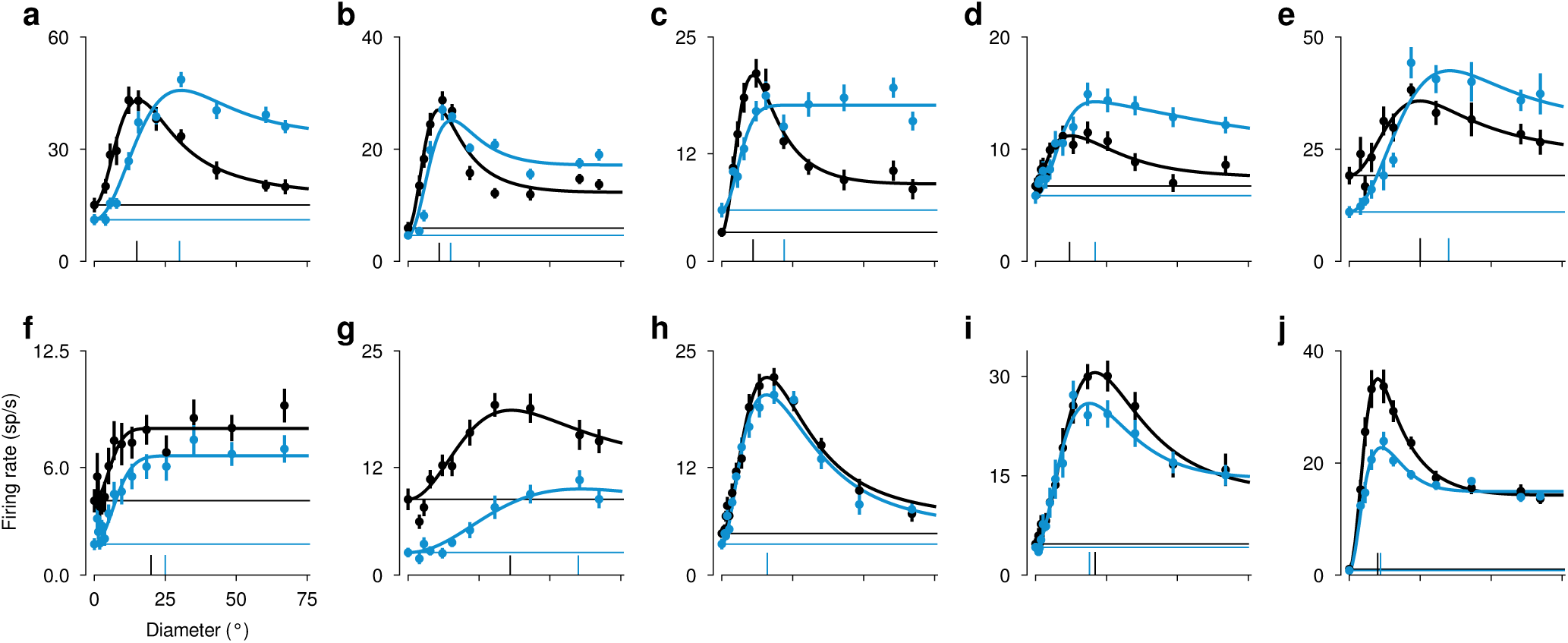
Size tuning curves of more dLGN example neurons. *Black*: control condition; *Blue*: V1 suppression; horizontal lines: responses to blank screen (size 0 deg); vertical lines: preferred size; error bars represent s.e.m.

**Figure S7.**
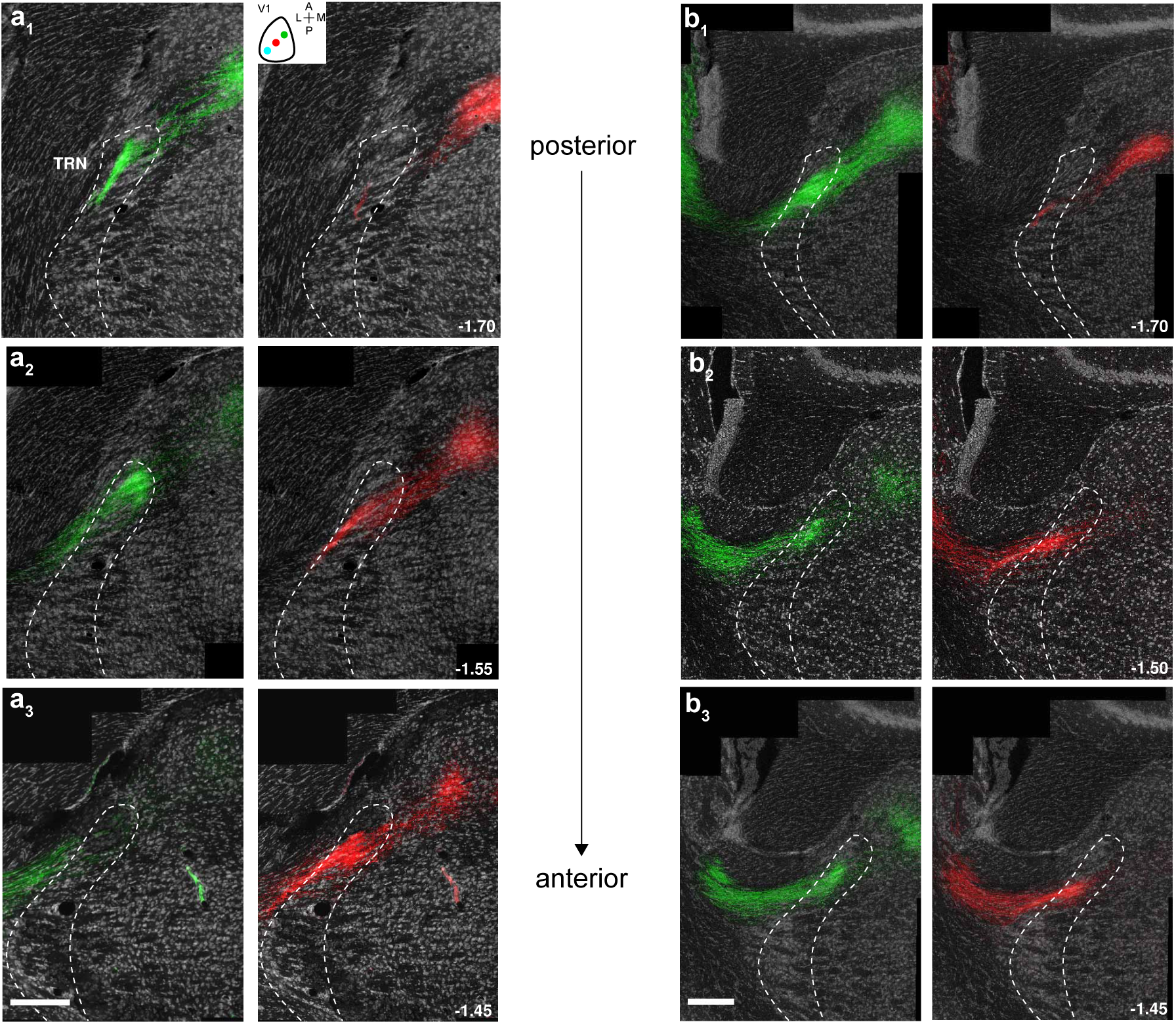
In TRN elevation is predominantly encoded along the anterior-posterior axis. (**a**_1_*_−_*_3_) Innervation pattern in TRN of axons from two L6CT populations transduced with pAAV-CAG-FLEX-EGFP (*green, left*) or pAAV-CAG-FLEX-mScarlet (*red, right*), respectively. V1 injections were performed along the retinotopic axis representing elevation (a_1_, *inset, right*), with EGFP labeling V1 regions representing higher elevations. Confocal images in (a_1*−*3_) are arranged from posterior to anterior (number indicates distance from bregma in mm); left and right images were taken from the same slice, with separate visualization of the two fluorophores. Note that more anterior regions in TRN contain terminal fields of L6CT axons labeled with mScarlet, i.e. representing lower elevations in the visual field; more posterior regions in TRN contain terminal fields of L6CT axons labeled with EGFP, i.e. representing higher elevations in the visual field. Scale bar 0.25 mm. (**b**_1_*_−_*_3_) Same as (a) for a second example mouse.

**Figure S8.**
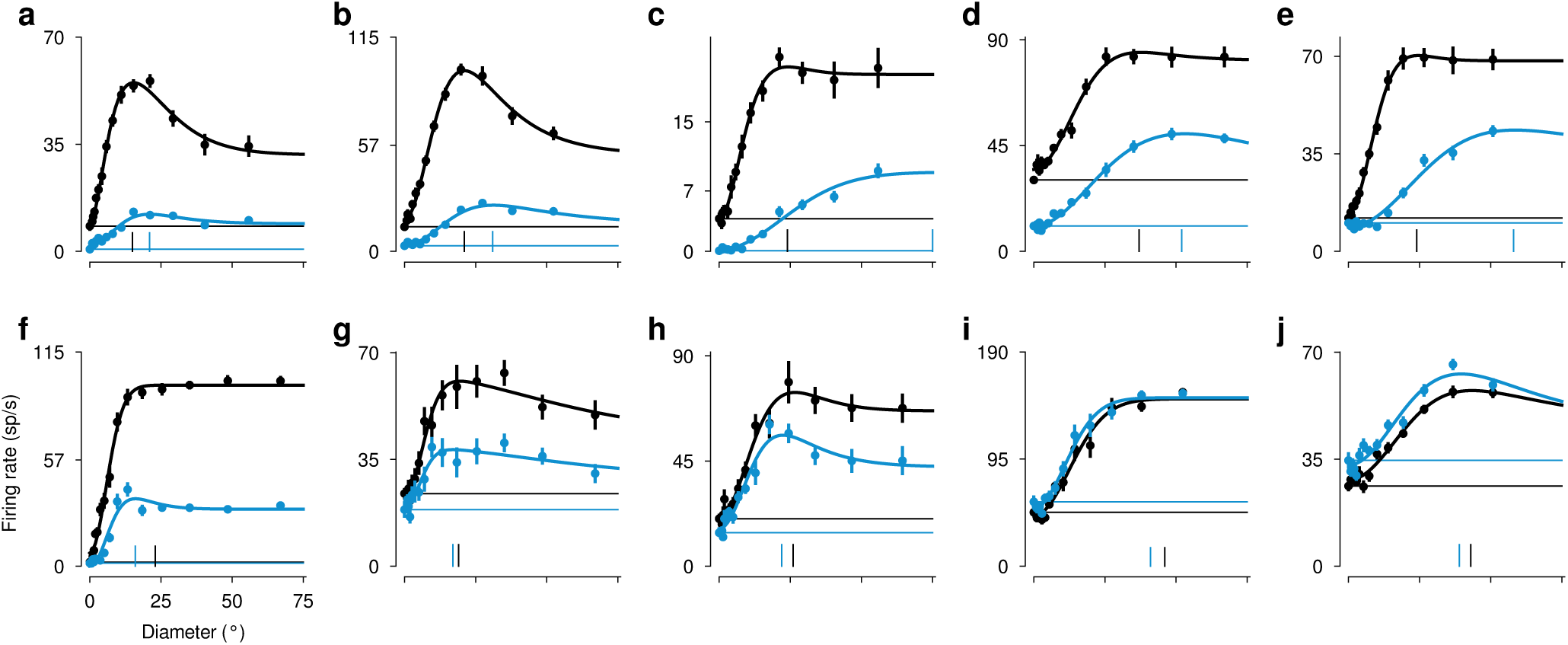
Size tuning curves of more visTRN example neurons. *Black*: control condition; *Blue*: V1 suppression; horizontal lines: responses to blank screen (size 0 deg); vertical lines: preferred size; error bars represent s.e.m.

**Figure S9.**
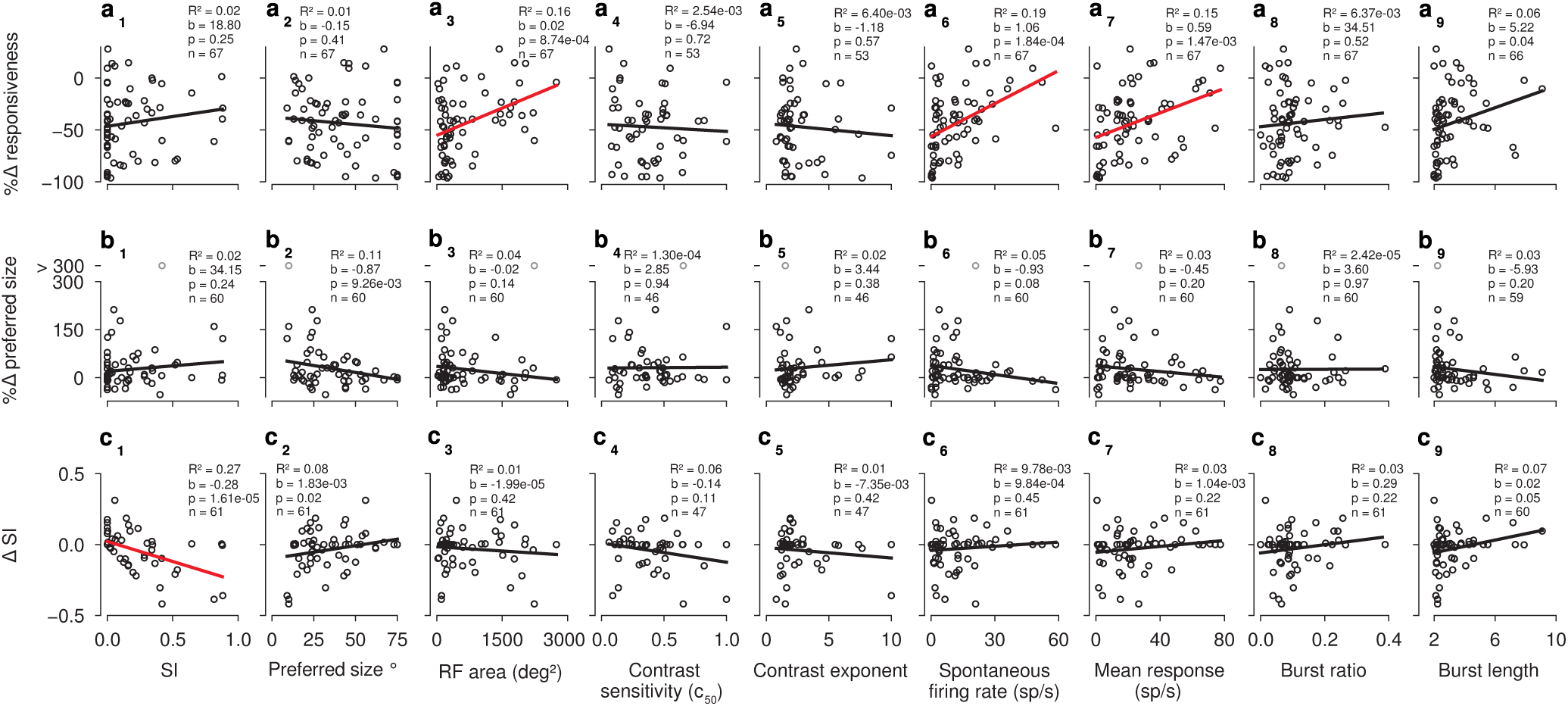
The relationship between CT feedback effects on visTRN neurons and their response properties. (**a**) Percent change in overall responsiveness by CT feedback as a function of SI (a_1_) and preferred size (a_2_) under control conditions, RF area as measured by a sparse noise stimulus (a_3_), contrast sensitivity (*c*_50_, a_4_) and steepness of the contrast response function (a_5_), spontaneous firing rate (a_6_), mean response (a_7_), burst ratio (a_8_), and burst length (a_9_) under control conditions of the size tuning experiments. While many relationships are not significant, CT feedback reduces overall responsiveness more for visTRN neurons with small compared to large RFs (a_3_), but the explained variance is small, partially because there is a wide array of effects for visTRN neurons with rather small RF coverage. Second, visTRN neurons with higher firing rates, show stronger CT feedback related modulations of firing rate (a_6_*_−_*_7_), pointing towards a multiplicative mechanism. (**b-c**) Same as (a), for CT feedback effects on preferred size and on surround suppression (SI), respectively. The observation that visTRN neurons with stronger surround suppression in control conditions show more pronounced changes in SI than those with weaker surround suppression (c_1_) could point towards an interesting subpopulation of visTRN neurons, which might represent spatial context and for which this representation is further enhanced by CT feedback. *Black*/*red line*: regression fit; *b*: slope; *p*: significance of slope.

**Figure S10.**
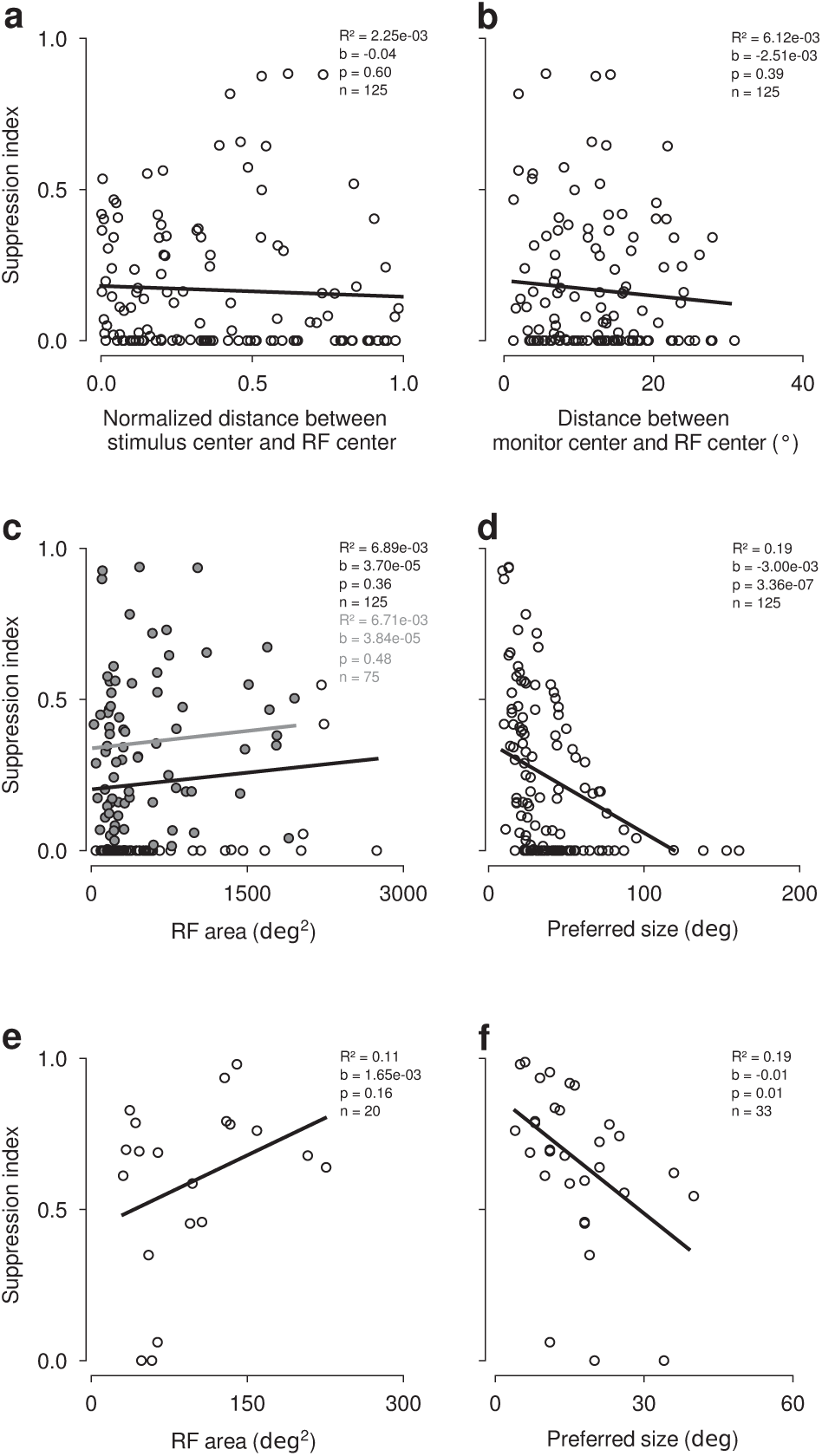
Correlations between suppression index and distance of RF center to monitor or stimulus center, and between suppression index and preferred size. (**a**) Suppression indices for visTRN population (*n* = 125) plotted against the normalized distance between stimulus center and their RF centers (*Black line*: linear regression; *b*: slope; *p*: signficance of slope). (**b**) Suppression indices for visTRN population plotted against the distance between monitor center and their RF centers. (**c-d**) Strength of surround suppression in visTRN measured during size tuning as a function of RF area mapped with the sparse noise stimulus (c) and as a function of preferred size taken from the size tuning curve (d). *Black*: regression line including all data points, *grey* regression line including a restricted set (*SI >* 0.01 and RF area *<* 2000 deg^2^). (**e-f**) Same as (c-d), for dLGN neurons. Note that in both visTRN (d) and dLGN (f) neurons with larger preferred sizes also tend to have less surround suppression. Although one caveat is the limited size of our monitor which for neurons with larger RFs might not allow for a sufficiently strong stimulation of the surround, this anti-correlation is in line with previous results from macaque V1^146^ and mouse V1^145^.

**Table S1.** eDOG parameters used to simulate dLGN size-tuning curves in Figure 3

|  |  |
| --- | --- |
| **integrator settings** |  |
| number of spatial points (Nr) | 2<sup>7</sup> |
| spatial resolution (dr) | 1 deg |
| number of temporal points (Nt) | 2<sup>1</sup> |
| temporal resolution (dt) | 1 ms |
| **stimulus settings** |  |
| spatial frequency | 0.05 cyc/deg |
| temporal frequency | 0 cyc/ms |
| patch diameter | start: 0 deg<br>stop: 75 deg<br>count: 76 |
| **ganglion difference-of-Gaussians** |  |
| center Gaussian | amplitude (A): 1<br>width (a): 3.33 deg |
| surround Gaussian | amplitude (B): 0.2<br>width (b): 7.67 deg |
| **spatial coupling kernels** |  |
| excitatory feedforward kernel (Krg) | weight (w): 1<br>amplitude (A): 1<br>width (a): 1 deg |
| inhibitory feedforward kernel (Krig) | weight (w): 0.5<br>amplitude (A): -1<br>width (a): 3 deg |
| excitatory feedback kernel (Krc_ex) | weight (w): 0 / 1<br>amplitude (A): 0.3<br>width (a): 1 deg |
| inhibitory feedback kernel (Krc_in) | weight (w): 0 / 1<br>amplitude (A): -0.6<br>width (a): 1 deg - 40 deg,<br>stepsize: 1 deg |

